# Early-Stage Corticostriatal Circuit Hyperactivity Impairs Cholinergic Function and Cognitive Flexibility in an Alzheimer’s Model

**DOI:** 10.64898/2026.01.13.699380

**Authors:** Yufei Huang, Xueyi Xie, Zhenbo Huang, Ruifeng Chen, Himanshu Gangal, Xuehua Wang, Karienn Souza, Julia Hunter, Xin Wu, Doodipala Samba Reddy, Jeannie Chin, Jun Wang

## Abstract

Cognitive flexibility deficits are a hallmark of early-stage Alzheimer’s disease (AD), but the underlying circuit mechanisms remain poorly understood. Here, we show that the 5xFAD mouse model of AD neuropathology exhibited early deficits in instrumental reversal learning, indicating cognitive inflexibility preceding spatial memory deficits. This impairment was associated with excessive neuronal reactivation in the medial prefrontal cortex (mPFC) and dorsomedial striatum (DMS), key regions for goal-directed behavior. Electrophysiological recordings revealed that mPFC neurons in young 5xFAD mice were hyperexcitable and received elevated excitatory input. Moreover, the mPFC-to-direct pathway medium spiny neuron (dMSN) circuit and the dMSNs themselves were selectively hyperactive. These hyperactive dMSNs exerted increased inhibitory control over cholinergic interneurons (CINs) in the DMS, coinciding with reduced CIN firing and diminished striatal acetylcholine (ACh) release. Critically, sustained chemogenetic inhibition of the mPFC-to-DMS circuit in 5xFAD mice reduced cortical Aβ accumulation, normalized glutamatergic transmission in both the mPFC and DMS, restored striatal ACh levels, and rescued reversal learning deficits. Together, these findings identify a hyperactive mPFC-to-DMS circuit that disrupts corticostriatal and cholinergic signaling, contributing to cognitive inflexibility in 5xFAD mice. Targeting this circuit may offer a therapeutic strategy to preserve cognitive function in the early stages of AD.

**Highlights:**

1. 5xFAD mice exhibit early cognitive deficits in instrumental reversal learning.
2. mPFC neurons and corticostriatal circuits are hyperactive, while cholinergic neurons are hypoactive in 5xFAD mice.
3. Sustained inhibition of mPFC-to-DMS circuit hyperactivity normalizes glutamatergic transmission and slows Aβ accumulation in 5xFAD mice.
4. Sustained inhibition of the mPFC-to-DMS circuit rescues reversal learning deficits in 5xFAD mice.

## INTRODUCTION

Alzheimer’s disease (AD) is the most common cause of dementia worldwide, affecting over 10% of individuals over 65 years old in the United States^1^. AD is characterized by the accumulation of amyloid-beta (Aβ) plaques and tau neurofibrillary tangles and by a progressive impairment of cognitive function^2–5^. Despite the development of FDA-approved Aβ-targeting antibodies and acetylcholinesterase (AChE) inhibitors, current treatments have limited effectiveness at later stages of the disease, when substantial neuronal loss has already occurred^6^. Therefore, identifying mechanisms driving early-stage pathophysiology is critical for developing therapeutic strategies capable of preserving cognitive function and modifying disease progression.

Although both Aβ and tau play critical roles in AD, there is evidence that the increase in Aβ occurs first and drives the preclinical and early stages of the disease^7–9^. Therefore, much effort has been devoted to understanding the mechanisms by which Aβ affects neuronal function early in AD. Several studies have demonstrated that Aβ accumulation disrupts glutamate reuptake, establishing a vicious cycle between Aβ deposition and neuronal hyperactivity^10–13^. Notably, Aβ accumulation initially emerges in cortical areas, including the medial prefrontal cortex (mPFC), and subsequently extends to innervated regions that include the hippocampus and striatum^14–16^. Although much attention has been paid to understanding how Aβ affects hippocampal function, less is known about how Aβ affects striatal circuits.

The dorsomedial striatum (DMS), a major target of mPFC projections, plays a critical role in goal-directed behavior and cognitive flexibility, which are core components of executive function that are compromised early in AD^17–25^. Cognitive flexibility refers to the ability of individuals to adapt their behaviors in response to changing circumstances or environmental cues^26^. Within the DMS, principal medium spiny neurons (MSNs) and cholinergic interneurons (CINs) interact to support this flexibility. CINs release acetylcholine (ACh), which regulates MSN activity to guide adaptive, goal-directed behaviors^27, 28^. Although cholinergic dysfunction is a hallmark of AD, and the striatum is one of the brain regions with the highest AChE activity^29^, the impact of Aβ on striatal cholinergic signaling is not well characterized. In particular, it remains unclear whether cortical Aβ disrupts CIN function in the DMS and whether such disruption contributes to early deficits in cognitive flexibility. Addressing these questions could clarify the circuit-level basis of early executive dysfunction in AD and reveal novel therapeutic targets to restore cognitive function and alter disease trajectory.

To address this knowledge gap, we used the 5xFAD mouse model of AD neuropathology^30^. We demonstrated that early cortical hyperactivity within the mPFC-to-DMS pathway disrupted cholinergic signaling in the striatum and contributed to reversal learning deficits. Sustained chemogenetic inhibition of this pathway attenuated Aβ accumulation, normalized corticostriatal excitatory transmission, restored cholinergic tone, and rescued cognitive flexibility. These findings reveal a circuit-level mechanism linking early Aβ pathology to executive dysfunction and suggest that targeting corticostriatal hyperactivity may offer a therapeutic entry point to slow disease progression in early-stage AD.

## RESULTS

### Three-month-old 5xFAD mice show instrumental reversal learning deficits indicative of cognitive inflexibility

We examined reversal learning in young 5xFAD mice to assess cognitive flexibility at the early stages of Aβ deposition. In this task (Fig. 1A), 3-month-old mice were trained to associate the left lever (A1) with grain pellets (O1) and the right lever (A2) with purified pellets (O2), as established in prior studies^23, 25, 31–33^. These two pellet types are similarly palatable and preferred over standard chow, with no intrinsic bias. To verify contingency learning, we performed a devaluation test in which mice were pre-fed either O1 or O2 prior to a choice test. Animals that had successfully learned the action-outcome association pressed the lever associated with the devalued (pre-fed) outcome less. After confirming initial learning, we reversed the action-outcome contingencies, such that A1 now delivered O2 and A2 delivered O1 (Reversal RR20 schedule; Fig. 1B). Successful adaptation to this reversal is widely used as a behavioral measure of cognitive flexibility^34^.

**Figure 1.**
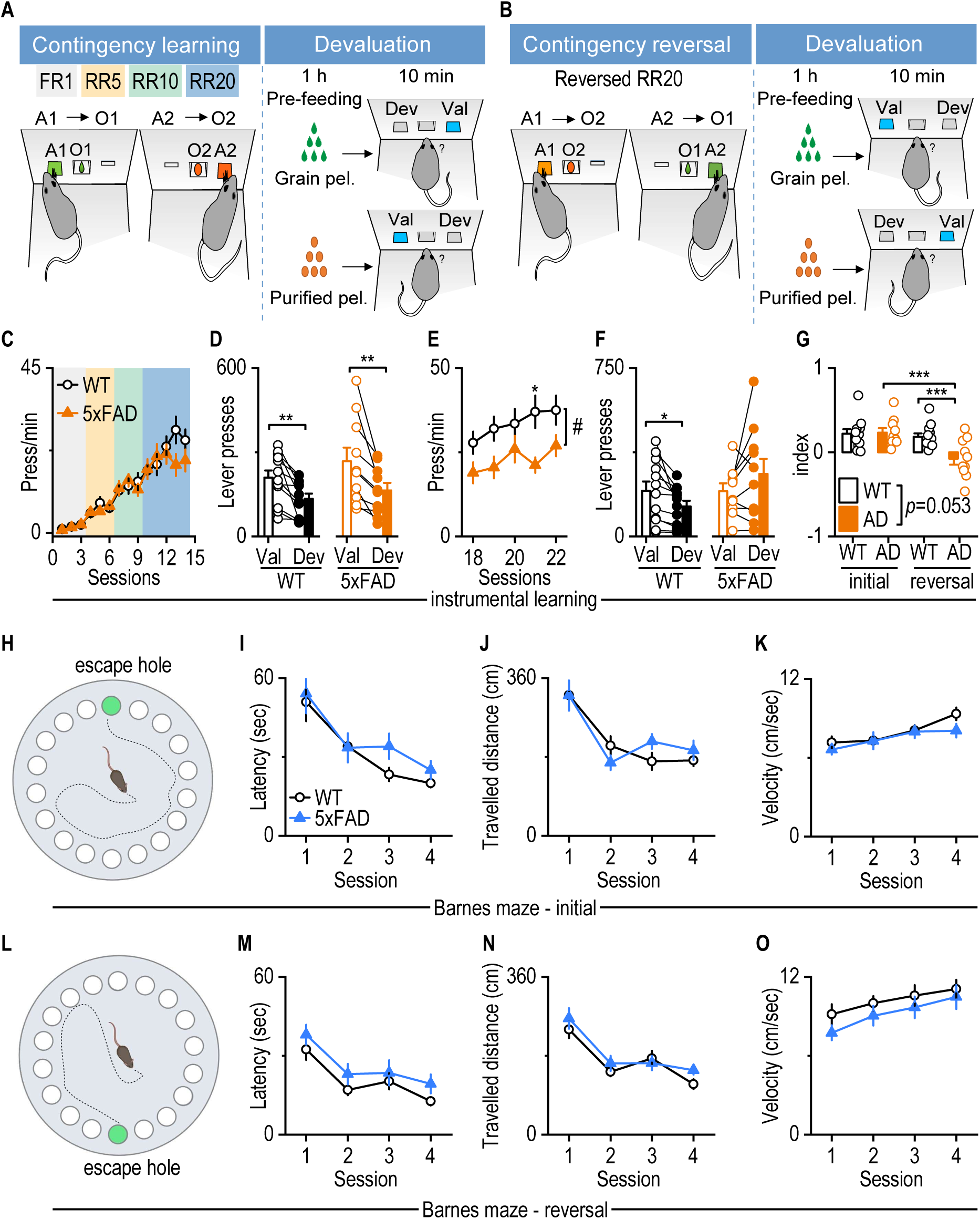
Early cognitive inflexibility emerges in 5xFAD mice during instrumental reversal learning. **A**, **B**, Schematic depiction of the instrumental conditioning and devaluation-test procedures for initial (A) and reversal (B) contingency learning. Mice initially learned to associate two actions of left or right lever pressing (A1 and A2) with receiving either the outcome of grain or purified pellets (O1 and O2). The learned contingencies were verified through devaluation tests, where mice were pre-fed with one outcome (O1 or O2) before a choice test between A1 and A2. Subsequently, mice underwent new learning with reversed contingencies, followed by a second round of devaluation and choice testing. FR1, fixed ratio 1, with one lever press for one reward delivery; RR5, random ratio 5, a probability of 0.2 for reward delivery; RR10, random ratio 10, a probability of 0.1 for reward delivery; RR20, a probability of 0.05 for reward delivery. **C**, 3-month-old wild-type (WT) and 5xFAD mice exhibited similar lever-press rates during the initial contingency learning. Two-way RM ANOVA with Greenhouse–Geisser correction followed by Sidak’s multiple comparisons test. n = 12 (WT) and 10 (5xFAD) mice. **D,** Both WT and 5xFAD mice displayed significantly fewer lever presses for devalued (Dev) than valued outcomes (Val) during the initial devaluation test. Paired t-test for WT and 5xFAD, \**p* < 0.05, \*\**p* < 0.01. n = 12 (WT) and 10 (5xFAD) mice. **E,** During reversal contingency training, 5xFAD mice exhibited lower pressing rates than WT mice. Mixed-effects model (REML), ^#^*p* < 0.05; Sidak’s multiple comparisons post-hoc analysis, \**p* < 0.05. n = 12 (WT) and 10 (5xFAD) mice. **F**, WT, but not 5xFAD, mice displayed significantly fewer lever presses for devalued (Dev) than valued outcomes (Val) during the reversal devaluation test. Paired t-test, \**p* < 0.05. n = 12 (WT) and 10 (5xFAD) mice. **G**, 5xFAD mice exhibited a significantly lower reversal index than their initial devaluation index as well as the reversal devaluation index of WT mice. The index is calculated as (Val-Dev)/(Val + Dev). Two-way RM ANOVA, *p* = 0.053; Sidak’s multiple comparisons post-hoc analysis, \*\*\**p* < 0.001. n = 12 (WT) and 10 (5xFAD, AD) mice. **H,** Schematic depiction of initial learning phase in Barnes maze. **I-K,** 3-month-old 5xFAD mice did not exhibit significant differences in latency (I), travel distance (J), and velocity (K) in finding the escape box compared with WT mice in the initial learning in the Barnes maze. Two-way RM ANOVA followed by Sidak’s multiple comparisons test. n = 22 (WT) and 16 (5xFAD) mice. **L,** Schematic depiction of reversal learning phase in Barnes maze. The escape hole was relocated opposite to the position of initial learning. **M-O,** 3-month-old 5xFAD mice did not exhibit significant differences in latency (M), travel distance(N), and velocity(O) in finding the escape box compared with WT mice in the reversal learning in the Barnes maze. Two-way RM ANOVA followed by Sidak’s multiple comparisons test for M and O. Generalized Linear Mixed Models for N. n = 22 (WT) and 16 (5xFAD) mice.

During the initial training phase, 3-month-old wild-type (WT) and 5xFAD mice showed similar lever-pressing rates (Fig. 1C, *F*_(1,20)_ = 0.4, *p* = 0.534). Both groups increased their responses in subsequent sessions (Fig. 1C, *F*_(1,13)_ = 46.649, *p* < 0.001), indicating intact initial learning. To evaluate whether both groups learned the action-outcome contingency, we performed the devaluation test described above. We found that both WT and 5xFAD mice pressed less on devalued levers (associated with the pre-fed outcome) than on valued levers (associated with the non-pre-fed outcome). This demonstrated successful learning of the initial action-outcome contingencies (Fig. 1D, *t*_11_ = 3.942, *p* = 0.0023 for WT; *t*_9_ = 3.628, *p* = 0.0055 for 5xFAD).

During the reversal RR20 learning phase, 5xFAD mice showed significantly reduced lever-pressing rates, as compared to WT mice (Fig. 1E, *F*_(1,20)_ = 5.434, *p* = 0.0303). Furthermore, while WT mice pressed the devalued lever less than the valued lever during the devaluation test, 5xFAD mice did not (Fig. 1F, *t*_11_ = 2.954, *p* = 0.0131 for WT; *t*_9_ = -1.551, *p* = 0.155 for 5xFAD). These results suggested that 5xFAD mice exhibited a deficit in acquiring the new action-outcome contingencies. Quantification of the devaluation index further confirmed this impairment: 5xFAD mice showed a significantly lower reversal learning index, as compared to their own initial devaluation performance (Fig. 1G, *F*_(1,20)_ = 4.216, *p* = 0.053; t = 4.49, *p* < 0.001). Additionally, the reversal devaluation index in 5xFAD mice was significantly lower than in WT controls (Fig. 1G, t = 0.341, *p* = 0.002), suggesting impaired cognitive flexibility during instrumental reversal learning.

To determine whether these deficits generalized to other cognitive domains, we assessed spatial memory using the Barnes maze (Fig. 1H and L)^35, 36^. At 3 months of age, 5xFAD and WT mice showed no significant differences in latency to locate the target escape chamber, distance traveled, or movement speed during both initial and reversal phases (Fig. 1I, *F*_(1, 36)_ = 0.9137, *p* = 0.3455; Fig. 1J, *F*_(1, 36)_ = 0.1402, *p* = 0.7103, Fig. 1K,, *F*_(1, 36)_ = 0.6623, *p* = 0.4211; Fig. 1M, *F*_(1, 36)_ = 2.184, *p* = 0.1482; Fig. 1N, *F*_(1, 36)_ = 1.382, *p* = 0.2476; Fig. 1O, *F*_(1, 36)_ = 1.117, *p* = 0.2975). This result indicated that 5xFAD mice had no spatial learning or memory impairments at this age.

We next examined whether the reversal learning deficit persisted at later stages by testing 6-month-old 5xFAD and WT mice using the same instrumental reversal task. During initial training, both genotypes showed similar lever-pressing rates (Supplementary Figure 1A), and both groups learned the initial action-outcome contingencies, although this was less robust in 5xFAD mice (Supplementary Figure 1B). Lever-pressing rates remained comparable in the reversal learning phase (Supplementary Figure 1C). However, while WT mice reduced their responses for the devalued outcome in the reversal devaluation test, 5xFAD mice did not (Supplementary Figure 1D), suggesting continued impairment in learning new action-outcome associations in 5xFAD mice. Devaluation index analysis confirmed a significant impairment in reversal learning in 5xFAD mice, compared to both their initial learning and WT mice (Supplementary Figure 1E). This indicated that cognitive flexibility deficits were still present in 6-month-old 5xFAD mice.

To assess spatial learning at 6 months of age, we repeated the Barnes maze test. In contrast to the 3-month-old cohort, 5xFAD mice at this age showed longer latencies and traveled greater distances to locate the escape hole during both the initial and reversal learning phases (Supplementary Figure 1F-I), indicating spatial learning deficits. Importantly, spontaneous locomotor activity was similar in WT and 5xFAD mice of both sexes (Supplementary Figure 2), excluding motor dysfunction as a confounding factor.

Together, these findings indicate that 5xFAD mice display cognitive inflexibility in instrumental reversal learning as early as 3 months of age, while spatial learning deficits emerge later, by 6 months of age.

### Reversal learning in 5xFAD mice reactivates excessive mPFC and DMS neurons

Given that 5xFAD mice exhibited intact initial learning and impaired reversal learning, we next examined whether this deficit was associated with an abnormal recruitment or reactivation of task-relevant neural ensembles. The mPFC and DMS represent key regions to investigate because they are critical for flexible, goal-directed behaviors^31–33, 37–42^. To monitor neuronal activation during instrumental learning, we employed the ArcTRAP system^43–45^. This system uses the immediate early gene, *Arc*, to label recently activated neurons. Arc is rapidly induced by experience-dependent activity and is well-established as a marker of neural engagement in learning and memory^46^. We crossed ArcTRAP mice with Ai14 and 5xFAD lines to generate ArcTRAP;Ai14;5xFAD and control (ArcTRAP;Ai14) cohorts.

To capture learning-related neuronal activation, 4-hydroxytamoxifen (4-OHT) was administered immediately after the second and third RR20 training sessions during both initial and reversal learning phases, triggering tdTomato expression in Arc-expressing neurons (Fig. 2A). We then quantified tdTomato-labeled neurons in the mPFC, DMS, and dentate gyrus of the hippocampus. During initial learning, ArcTRAP;Ai14 controls and ArcTRAP;Ai14;5xFAD mice showed comparable densities of activated neurons in the mPFC and DMS (Fig. 2B, *t*_11_ = 1.499, *p* = 0.5106; Fig. 2C, *U* = 13, *p* = 0.2949). However, during reversal learning, ArcTRAP;Ai14;5xFAD mice displayed significantly greater neuronal activation than ArcTRAP;Ai14 control mice. This difference was significant in both the mPFC (Fig. 2D, *U* = 0, *p* = 0.0159) and DMS (Fig. 2E, *t*_3.389_ = 3.609, *p* = 0.0299). In contrast, there was no significant difference in the number of activated neurons in the hippocampal dentate gyrus between learning phases in either ArcTRAP;Ai14 or ArcTRAP;Ai14;5xFAD mice (Supplementary Figure 3). These findings suggest that the mPFC and DMS, but not the hippocampus, are more active in 5xFAD animals than in controls during reversal learning.

**Figure 2.**
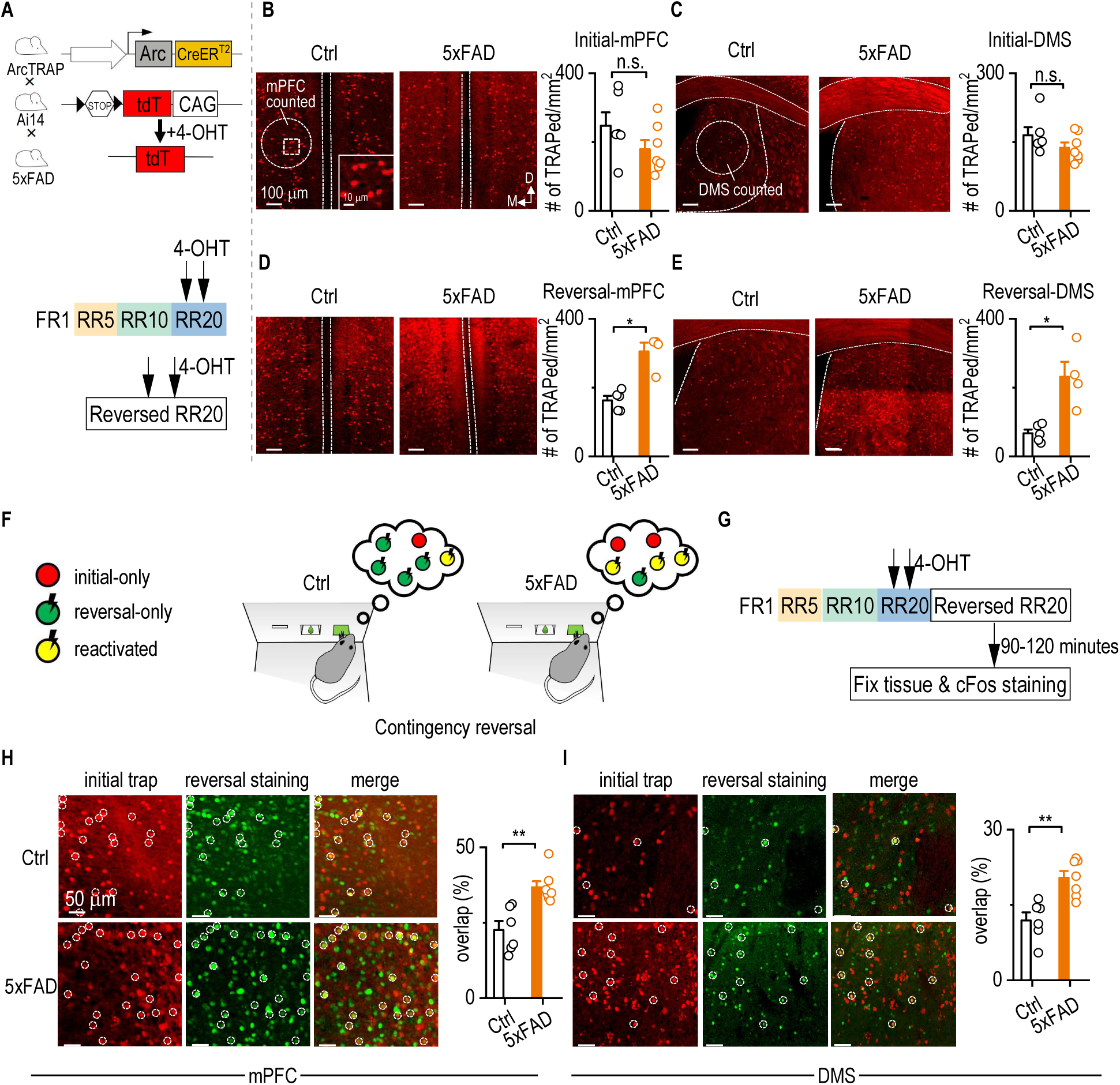
Reversal learning in 5xFAD mice reactivates excessive mPFC and DMS neurons. **A,** Schematic diagrams illustrating the ArcTRAP mechanism and experimental trapping strategy. 4-hydroxytamoxifen (4-OHT) was administered to induce tdTomato expression, labeling neurons activated during either the middle two sessions of initial learning or reversal learning under an RR20 schedule. **B,** Representative images of tdTomato-labeled mPFC neurons activated during initial learning in control (Ctrl) and 5xFAD mice. No significant difference in DMS activation was observed between groups. Unpaired t-test. n = 6 mice for Ctrl and 7 mice for 5xFAD. **C,** Representative images of tdTomato-labeled DMS neurons activated during initial learning in Ctrl and 5xFAD mice. No significant difference in DMS neuron activation was observed between groups. Mann-Whitney test. n = 6 mice for Ctrl and 7 mice for 5xFAD. **D,** Representative images of mPFC neurons activated during reversal learning. 5xFAD mice showed increased mPFC neuron activation compared to controls. Mann-Whitney test, \**p* < 0.05. n = 5 mice for Ctrl; 4 mice for 5xFAD. **E,** Representative images of DMS neurons activated during reversal learning. 5xFAD mice exhibited greater DMS neuron activation than Ctrl mice. Unpaired t test with Welch’s correction, \**p* < 0.05. n = 5 mice for Ctrl; 4 mice for 5xFAD. **F and G,** Schematic illustration of longitudinal tracking of neuronal activation during reversal learning. Neurons activated during initial learning were permanently labeled with tdTomato via the ArcTRAP strategy. After reversal learning, c-Fos staining labeled newly activated neurons with GFP. Neurons active in both phases co-expressed tdTomato and GFP. **H,** Representative images of mPFC neurons co-labeled with tdTomato and GFP in Ctrl and 5xFAD mice. 5xFAD mice displayed a significantly higher overlap between initial- and reversal-learning activated mPFC neurons. Mann-Whitney Rank Sum Test, \*\**p* < 0.01. n = 6 mice for Ctrl; 7 mice for 5xFAD. **I,** Representative images of DMS neurons co-labeled with tdTomato and GFP. 5xFAD mice exhibited greater reactivation of DMS neurons during reversal learning compared to Ctrl mice. Mann-Whitney Rank Sum Test, \*\**p* < 0.01. n = 6 mice for Ctrl; 7 mice for 5xFAD.

To determine whether this heightened activation reflected aberrant re-engagement of the same neuronal ensembles involved in initial learning, we performed a longitudinal tagging experiment. Neurons activated during initial learning were labeled using tdTomato (via ArcTRAP;Ai14), while those recruited during reversal learning were identified using c-Fos immunostaining (Fig. 2F and 2G). This approach allowed us to quantify the proportion of initial learning-activated neurons (tdTomato+) that were reactivated during reversal learning (tdTomato+ & c-Fos+) within the same animal. We found that ArcTRAP;Ai14;5xFAD mice exhibited significantly greater co-expression of tdTomato and c-Fos in mPFC neurons (Fig. 2H, *U* = 0, *p* = 0.0012) and DMS neurons (Fig. 2I, U = 1, *p* = 0.0023), as compared to ArcTRAP;Ai14 mice.

### mPFC neurons in young 5xFAD mice are hyperexcitable and receive increased excitatory input

To examine whether functional abnormalities in mPFC neurons are also evident at the cellular and synaptic level, we next performed whole-cell recordings. We focused on 4-month-old 5xFAD mice because early-stage cortical amyloid pathology^14, 47^ and neuronal hyperactivity^12, 48^ has been reported at this age.

We performed whole-cell patch-clamp recordings from pyramidal neurons in the mPFC to assess their intrinsic excitability (Fig. 3A). Compared to age-matched WT controls, mPFC neurons from 5xFAD mice exhibited significantly higher firing rates in response to increasing current injections (Fig. 3B, *F*_(1,29)_ = 13.10, *p* = 0.0011), indicating greater intrinsic excitability. To examine synaptic input, we recorded spontaneous excitatory postsynaptic currents (EPSCs) (Fig. 3C). Both the amplitude (Fig. 3D, *U* = 20, *p* = 0.0199) and frequency (Fig. 3E, *t*_17.80_ = 2.196, *p* = 0.0416) of spontaneous EPSCs were significantly elevated in 5xFAD mice, suggesting enhanced baseline glutamatergic transmission.

**Figure 3.**
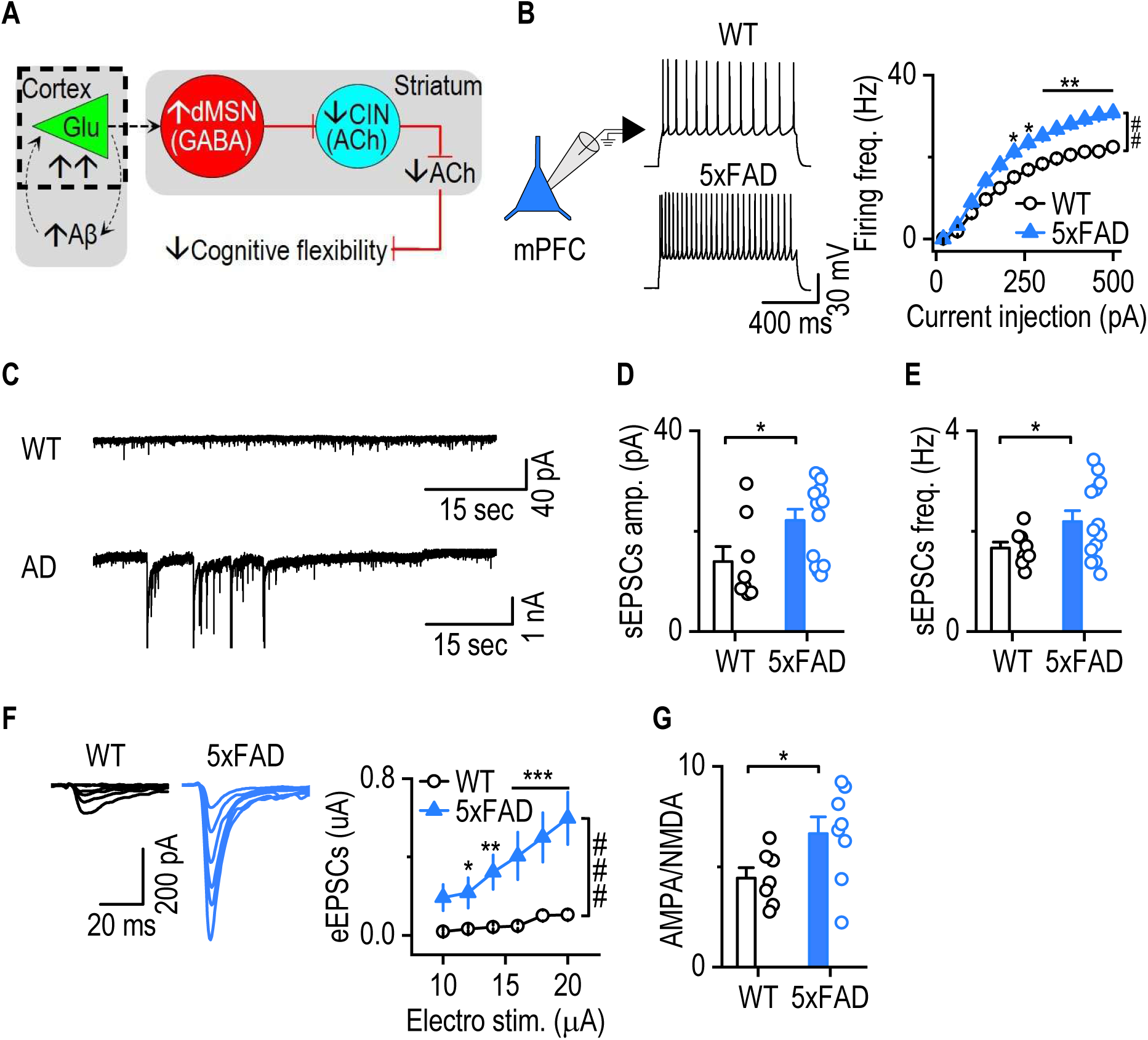
mPFC neurons in young 5xFAD mice are hyperexcitable and receive increased excitatory input. Coronal mPFC slices were prepared from 4-month-old WT and 5xFAD mice. **A,** Schematic of the proposed circuit mechanism underlying impaired cognitive flexibility in early-stage AD. Amyloid-β accumulation in the cortex drives hyperactivity of cortical neurons, particularly in the mPFC. This results in excessive glutamatergic input to direct-pathway MSNs (dMSNs), which in turn provide increased GABAergic inhibition to striatal cholinergic interneurons (CINs), impairing cholinergic signaling and cognitive flexibility. The dashed area indicates the experimental focus on hyperactivity of mPFC neurons in early AD. **B**, mPFC neurons had a significantly higher frequency of evoked firing in 5xFAD animals than WT controls. Two-way RM ANOVA with Greenhouse–Geisser correction, ^##^*p* < 0.01; Sidak’s multiple comparisons post-hoc analysis, \**p* < 0.05, \*\**p* < 0.01 versus WT at the same stimulating intensities. Current injection: 300 pA, 1 sec. n = 15 neurons from 3 mice (WT) and 16 neurons from 5 mice (5xFAD). **C**, Representative traces of spontaneous excitatory postsynaptic currents (sEPSCs) recorded from mPFC neurons in WT and 5xFAD mice. **D and E**, sEPSC amplitude (D) and frequency (E) were significantly increased in 5xFAD mice compared to WT. Mann-Whitney Rank Sum Test for amplitude; Unpaired t-test with Welch’s correction for frequency, \**p* < 0.05. n = 8 neurons from 3 mice (WT) and 13 neurons from 3 mice (5xFAD). **F**, Electrically evoked EPSCs (eEPSCs) in mPFC neurons were significantly larger in 5xFAD mice than WT controls. Two-way RM ANOVA with Greenhouse–Geisser correction, ^###^*p* < 0.001; Sidak’s multiple comparisons post-hoc analysis, \**p* < 0.05, \*\**p* < 0.01, \*\*\**p* < 0.001 versus WT at the same stimulating intensities. n = 13 neurons from 3 mice (WT) and 8 neurons from 3 mice (5xFAD). **G**, The AMPA/NMDA ratio was higher in 5xFAD mice than WT controls. Unpaired t-test, \**p* = 0.051. n = 7 neurons from 3 mice (WT) and 8 neurons from 3 mice (5xFAD).

We next evaluated evoked synaptic responses to assess functional synaptic drive. Electrically evoked EPSCs were significantly larger in 5xFAD mice than in WT controls (Fig. 3F *F*_(1,19)_ = 15.881, *p* < 0.001). The AMPA/NMDA ratio was also elevated in 5xFAD mice (Fig. 3G, *t*_13_ = -2.160, *p* = 0.0501), indicating a greater contribution of AMPA receptor-mediated currents.

Together, these findings demonstrate that mPFC neurons in young 5xFAD mice are intrinsically hyperexcitable and receive augmented excitatory synaptic input.

### The mPFC-to-dMSN circuit and dMSNs are selectively hyperactive in 5xFAD mice

Given the hyperexcitability of mPFC neurons observed in 5xFAD mice, we next investigated whether this elevated cortical activity altered downstream striatal function, particularly within the mPFC-to-DMS circuit. The mPFC projects directly to the DMS^49, 50^, a critical region for flexible behavior and reversal learning^21, 33^. We first examined excitatory synaptic transmission onto striatal MSNs, the principal output neurons of the DMS (Fig. 4A). Electrically evoked EPSCs were significantly larger in 5xFAD mice than in WT controls (Fig. 4B *F*_(1,40)_ = 6.836, *p* = 0.013), indicating enhanced excitatory transmission. In addition, spontaneous EPSC amplitudes were significantly higher in 5xFAD mice (Fig. 4C, *U = 43*, *p* = 0.0009), while the frequency remained unchanged (Fig. 4D *U = 85.5*, *p* = 0.1120). This suggested that the observed increase in synaptic transmission was primarily driven by postsynaptic changes rather than increased presynaptic activity.

**Figure 4.**
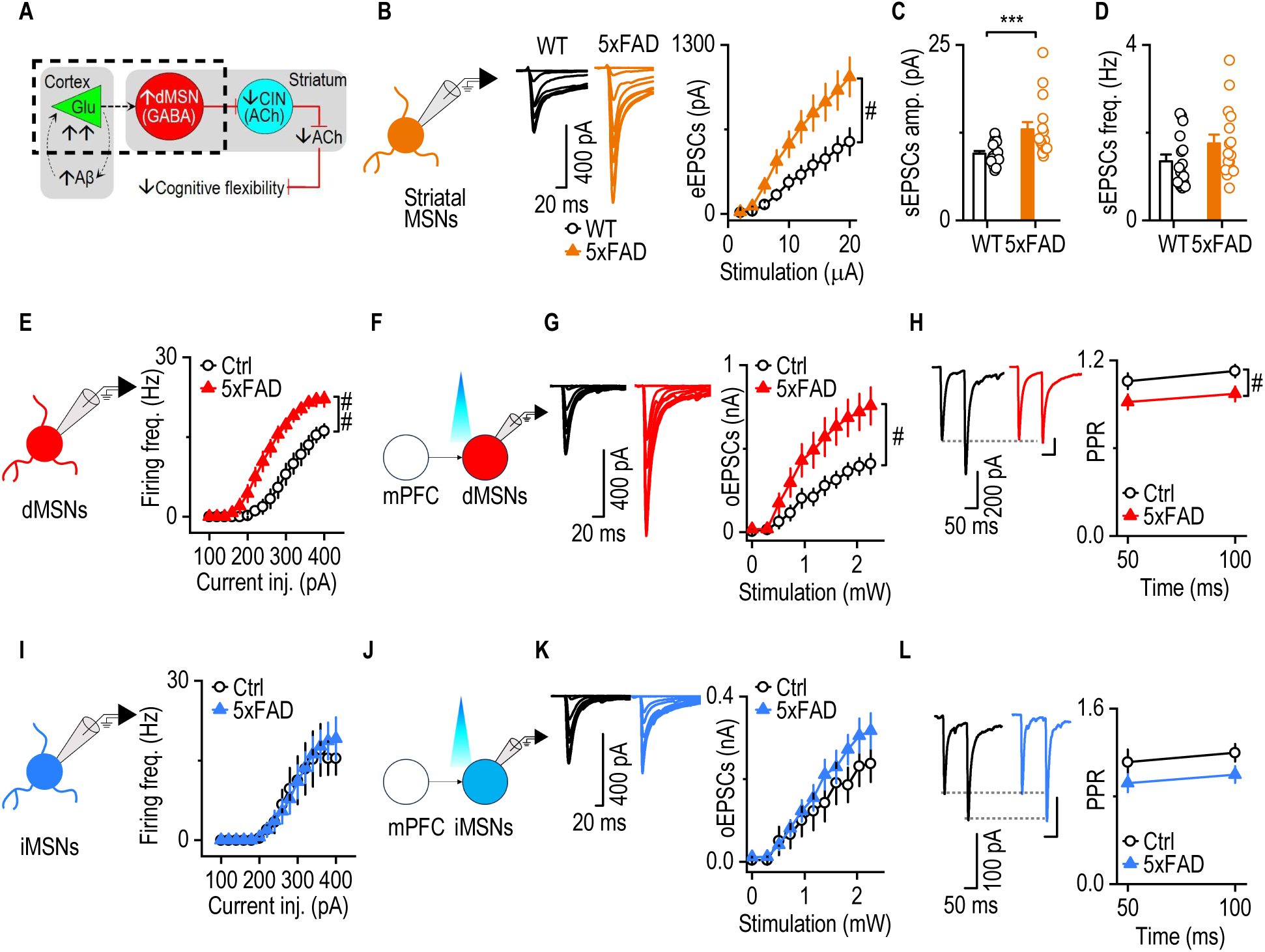
The mPFC-to-dMSN circuit and dMSNs are selectively hyperactive in 5xFAD mice. **A,** Schematic of the neurocircuit mechanism underlying decreased cognitive inflexibility in the early stage of AD. The dashed area indicates testing whether the mPFC-to-dMSN projection is hyperactive in early-stage AD. **B**, eEPSCs amplitude in DMS MSNs was greater in 4-month-old 5xFAD mice than in the WT control. Two-way RM ANOVA with Greenhouse–Geisser correction, ^#^*p* < 0.05; Sidak’s multiple comparisons post-hoc analysis, \**p* < 0.05, \*\**p* < 0.01, \*\*\**p* < 0.001 versus WT at the same stimulating intensities. n = 22 neurons from 5 mice (WT); 20 neurons from 5 mice (5xFAD). **C** and **D**, sEPSCs amplitude (C), but not frequency (D), was higher in 5xFAD mice than WT controls. Mann-Whitney U test, \**p* < 0.05. n = 16 neurons from 5 mice (WT and 5xFAD). **E,** Diagram illustrating patch clamp recording of dMSNs. dMSNs in 5xFAD mice had a higher evoked firing frequency than in Ctrl mice. Generalized Linear Mixed Models, ^##^*p* < 0.01. n = 8 neurons from 4 mice (Ctrl and 5xFAD) **F,** Diagram illustrating patch clamp recording of optical stimulation of mPFC inputs to dMSNs. **G,** oEPSCs amplitude in DMS dMSNs was greater in 4-month-old 5xFAD mice than in D1-tdTomato controls. Two-way RM ANOVA with Greenhouse–Geisser correction, ^#^*p* < 0.05. n = 10 neurons from 4 mice (WT); 15 neurons from 4 mice (5xFAD). **H**, PPR was smaller in 5xFAD mice than in controls. Two-way RM ANOVA, ^#^*p* < 0.05. n = 8 neurons from 4 mice (WT); 10 neurons from 4 mice (5xFAD). **I,** Diagram illustrating patch clamp recording of iMSNs. iMSNs in 5xFAD mice did not show a higher evoke firing frequency than in WT mice. Generalized Linear Mixed Models. n = 7 neurons from 4 mice (Ctrl); 9 neurons from 4 mice (5xFAD). **J,** Diagram illustrating patch clamp recording of optical stimulation of mPFC inputs to iMSNs. **K,** oEPSCs amplitude in DMS iMSNs was not significantly greater in 4-month-old 5xFAD mice than in WT controls. Two-way RM ANOVA with Greenhouse–Geisser correction. n = 10 neurons from 4 mice (Ctrl and 5xFAD). **L**, PPR of oEPSCs had no difference in 5xFAD mice than WT controls. Two-way RM ANOVA. N = 7 neurons from 4 mice (Ctrl and 5xFAD).

Since MSNs are subdivided into dMSNs and indirect-pathway MSNs (iMSNs), we next asked whether altered mPFC inputs affect one or both MSN subpopulations. To selectively activate mPFC projections, we infused AAV-Chronos-GFP into the mPFC of D1-tdTomato;5xFAD and D1-tdTomato (control) mice, enabling optogenetic stimulation of mPFC terminals in the DMS while distinguishing dMSNs (tdTomato-positive) from iMSNs (tdTomato-negative) (Supplementary Fig. 4A). In 4-month-old D1-tdTomato;5xFAD mice, dMSNs exhibited significantly greater excitability in response to current injections than they did in D1-tdTomato controls (Fig. 4E, *F*_(1, 14)_ = 11.03, *p* = 0.0050). Moreover, optogenetically evoked EPSCs in dMSNs were also significantly larger in D1-tdTomato;5xFAD mice (Fig. 4F and G, *F*_(1, 23)_ = 4.694, *p* = 0.0409), and the paired-pulse ratio (PPR) of the EPSCs was reduced (Fig. 4H, F_(1, 16)_ = 5.280, *p* = 0.0354), indicating increased presynaptic glutamate release from mPFC terminals. Notably, this hyperactivity persisted in 12-month-old D1-tdTomato;5xFAD mice, suggesting a sustained and progressive synaptic alteration (Supplementary Figure 4B and 4C).

In contrast, the mPFC-to-iMSN pathway appeared unaffected. Comparison of 4-month-old D1-tdTomato;5xFAD and D1-tdTomato control mice revealed no significant differences in iMSN excitability (Fig. 4I, *F*_(1, 14)_ = 0.2383, p = 0.6330), EPSC amplitude (Fig. 4J and K, *F*_(1, 18)_ = 0.8385, *p* = 0.3719), or PPR (Fig. 4L, F_(1, 12)_ = 2.715, *p* = 0.1253). In addition, no differences in these parameters were observed at 12 months of age (Supplementary Figure 4D and 4E). These *ex vivo* findings indicated that the observed hyperactivity was selectively associated with the mPFC-to-dMSN circuit rather than the mPFC-to-iMSN pathway.

To assess whether neuronal hyperactivity in 5xFAD mice is also evident at the network level in vivo, we examined seizure susceptibility and EEG activity. Using 6-Hz corneal stimulation (Supplementary Fig. 5A), we found that 5xFAD mice had a lower threshold for seizure induction (Supplementary Fig. 5B) and displayed significantly longer seizures of greater severity, as compared to WT mice (Supplementary Fig. 5C-D). We further confirmed this *in vivo* hyperactivity using EEG recordings before and after pilocarpine injection (Supplementary Fig. 5E). Following pilocarpine administration, 5xFAD mice exhibited significantly elevated spike rates relative to WT controls (Supplementary Fig. 5F, G). Survival analysis also showed a stark contrast between groups because 5xFAD mice failed to survive pilocarpine injection (Supplementary Fig. 5H), indicating increased seizure vulnerability.

Together, our patch-clamp recordings suggest that the mPFC–D1MSN circuit is hyperactive in 5xFAD mice, while our in vivo assays indicate that 5xFAD mice exhibit heightened neuronal excitability and seizure susceptibility at the network level.

### Enhanced dMSN-to-CIN inhibition coincides with reduced CIN activity and striatal ACh release in 5xFAD mice

Having demonstrated that dMSNs were selectively hyperactive in 5xFAD mice and given prior reports that dMSNs directly inhibited CINs in the striatum^25, 51^, we hypothesized that elevated dMSN activity would disrupt CIN function (Fig. 5A). To test this hypothesis, we optogenetically activated dMSNs while recording from CINs in D1-Cre;ChAT-eGFP;5xFAD and D1-Cre;ChAT-eGFP (control) mice (Fig. 5B). AAV-FLEX-Chrimson-tdTomato was infused into the DMS to selectively express Chrimson in dMSNs, and 590-nm light was used to activate these projections. CINs in D1-Cre;ChAT-eGFP;5xFAD mice exhibited significantly larger optically evoked inhibitory postsynaptic currents (IPSCs), as compared to D1-Cre;ChAT-eGFP controls (Fig. 5C, 5D; *F*_(1, 45)_ = 10.02, *p* = 0.0028). This indicated that dMSN-to-CIN inhibition was significantly enhanced in this mouse model of AD.

**Figure 5.**
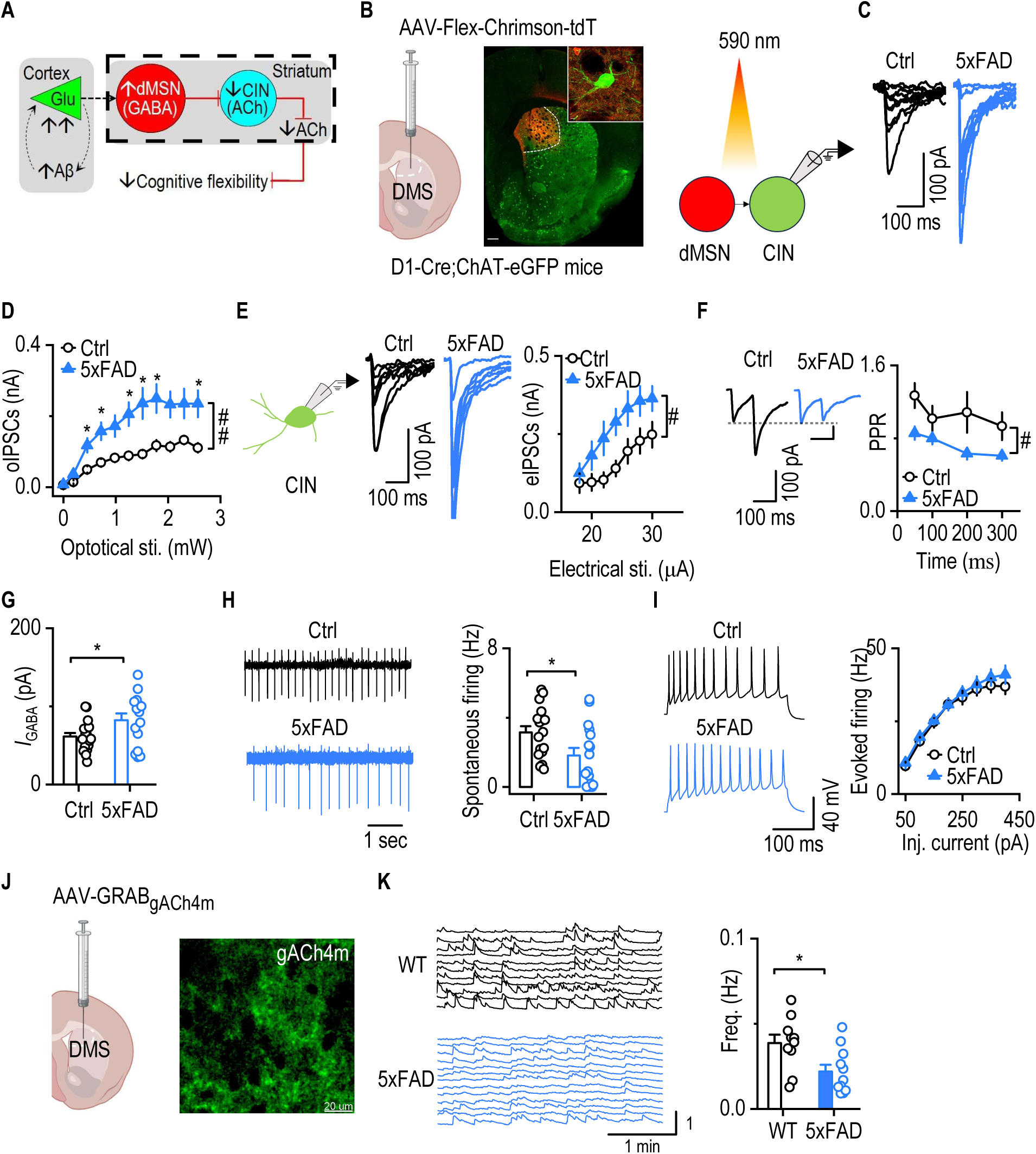
Enhanced dMSN-to-CIN inhibition coincides with reduced CIN activity and striatal ACh release in 5xFAD mice. CINs in the DMS were identified by GFP expression in brain slices from 5xFAD;ChAT-eGFP (5xFAD) and ChAT-eGFP (Ctrl) mice. **A,** Schematic illustration of the proposed circuit mechanism underlying impaired cognitive flexibility in early-stage AD. The dashed area highlights the experimental focus on changes in CIN physiology and dMSN-to-CIN synaptic transmission. **B,** Diagram showing viral infusion into the DMS and representative image of virus expression in D1-Cre;ChAT-eGFP mice. Right: schematic of oIPSCs recorded from CINs following dMSN stimulation. **C**, Representative oIPSC traces from dMSN-to-CIN synapses in Ctrl and 5xFAD mice. **D,** CINs in 5xFAD mice received significantly stronger oIPSCs from dMSNs compared to controls. Generalized Linear Mixed Model (GLMM), ^##^*p* < 0.01; Sidak’s multiple comparisons post-hoc analysis, \**p* < 0.05 WT versus 5xFAD at the same optical stimulation. n = 23 neurons from 3 mice (Ctrl); 24 neurons from 3 mice (5xFAD). **E**, Schematic of whole-cell patch-clamp recordings from CINs. CINs in 5xFAD mice displayed stronger electrically evoked IPSCs than those in Ctrl mice. Two-way RM ANOVA with Greenhouse–Geisser correction, ^#^*p* < 0.05. n= 14 neurons from 4 mice (Ctrl); 15 neurons from 4 mice (5xFAD). **F,** PPR was smaller in 5xFAD mice than in controls. Two-way RM ANOVA with Greenhouse–Geisser correction, ^#^*p* < 0.05. n= 14 neurons from 4 mice (Ctrl); 15 neurons from 4 mice (5xFAD). **G,** CINs in 5xFAD mice showed higher GABA (50 µM)-induced currents compared to Ctrl animals. Unpaired t-test with Welch’s correction, \**p* < 0.05. n= 18 neurons from 4 mice (Ctrl); 15 neurons from 4 mice (5xFAD). **H,** CINs in 5xFAD mice displayed reduced spontaneous firing rates compared to controls. Mann-Whitney Rank Sum Test, \**p* < 0.05. n= 17 neurons from 4 mice (Ctrl); 16 neurons from 4 mice (5xFAD). **I,** Evoked firing of CINs did not differ between the two groups of animals. Two-way RM ANOVA with Greenhouse–Geisser correction. n= 17 neurons from 4 mice (Ctrl); 14 neurons from 5 mice (5xFAD). **J,** Confocal image of DMS slices showing expression of the genetically encoded acetylcholine sensor gACh4m. **K,** Representative traces of spontaneous ACh release events in acute DMS slices from WT and 5xFAD mice. 5xFAD mice showed a significantly reduced frequency of spontaneous ACh release. unpaired t-test, \**p* < 0.05. n = 10 slices of 4 mice (WT); 11 slices of 4 mice (5xFAD).

Next, we investigated whether the enhanced dMSN-to-CIN inhibition observed with optogenetic activation could also be detected using electrical stimulation, which recruits all synaptic inputs, including those from dMSNs. To do so, we performed whole-cell recordings of IPSCs evoked by electrical stimulation. CINs from AD mice exhibited significantly larger IPSC amplitudes than WT mice (Fig. 5E; *F*_(1, 27)_ = 5.502, *p* = 0.0266). We then explored the mechanism underlying this increased CIN inhibition by evaluating presynaptic and postsynaptic components. PPR values were significantly reduced in D1-Cre;ChAT-eGFP;5xFAD mice (Fig. 5F; F_(1, 27)_ = 6.545, *p* = 0.0164), indicating increased presynaptic GABA release from inhibitory inputs onto CINs. To assess postsynaptic changes, we bath-applied GABA (50 µm) and measured the induced current in CINs. GABA-evoked responses were significantly greater in D1-Cre;ChAT-eGFP;5xFAD mice than in D1-Cre;ChAT-eGFP control mice (Fig. 5G; *t*_21.57_ = 2.155, *p* = 0.0426), indicating increased postsynaptic GABA_A_ receptor responsiveness. We then assessed the functional impact of this increased inhibitory tone on CIN activity. Spontaneous firing of CINs was significantly reduced in D1-Cre;ChAT-eGFP;5xFAD mice, as compared to D1-Cre;ChAT-eGFP controls (Fig. 5H; U = 72, *p* = 0.022), whereas evoked firing responses remained intact (Fig. 5I; F_(1,29)_ = 0.434, *p* = 0.515). This suggested that CIN hypoactivity was likely due to excessive inhibition rather than to changes in intrinsic excitability.

To assess whether reduced CIN activity translated to decreased cholinergic output, we expressed the genetically encoded ACh sensor, gACh4m^52^, in the DMS (Fig. 5J) of D1-Cre;ChAT-eGFP;5xFAD and D1-Cre;ChAT-eGFP control mice. We observed a marked reduction in the frequency of spontaneous ACh release events in D1-Cre;ChAT-eGFP;5xFAD mice, as compared to controls (Fig. 5K; t_19_ = 2.656, *p* = 0.0156), indicating diminished cholinergic tone. Given that the ACh released by CINs modulates MSNs indirectly via GABAergic interneurons^27^, we next examined whether CIN-to-MSN inhibition was also altered. To address this, we generated ChAT-Cre;Ai32;5xFAD and ChAT-Cre;Ai32 (control) mice, and optogenetically stimulated CINs while recording IPSCs in MSNs. We found that CIN-to-MSN IPSCs were significantly lower in ChAT-Cre;Ai32;5xFAD mice than in ChAT-Cre;Ai32 controls (Supplementary Figure 6).

Together, these findings suggested that enhanced dMSN-to-CIN inhibition in the 5xFAD model of AD was associated with reduced CIN activity and less striatal ACh release. This disruption in cholinergic signaling may contribute to the early cognitive inflexibility observed in AD.

### Sustained mPFC-to-DMS inhibition slows Aβ accumulation, reduces glutamatergic transmission, and enhances striatal ACh in 5xFAD mice

Given that mPFC hyperactivity emerges early in 5xFAD mice alongside cortical Aβ accumulation (Fig. 3) and that the mPFC-to-DMS circuit shows excessive output (Fig. 4) and is linked to downstream cholinergic suppression (Fig. 5), we next asked whether sustained inhibition of this pathway affected these pathophysiological changes. Prior studies have suggested the existence of a feed-forward loop in which neuronal hyperactivity promotes Aβ accumulation, which in turn impairs glutamate clearance and further increases neuronal excitability^13, 53, 54^. We, therefore, tested whether chemogenetic inhibition of the mPFC-to-DMS projection could reduce cortical Aβ accumulation, normalize glutamatergic output, and restore striatal cholinergic tone.

As a first step, we confirmed that acute chemogenetic inhibition of mPFC neurons reduced both intrinsic excitability and synaptic transmission in *ex vivo* slices (Supplementary Fig. 7). However, previous work indicated that sustained neuronal silencing over several weeks was necessary to impact Aβ pathology^55^. To implement a long-term circuit-specific inhibition strategy, we infused either AAV-DIO-hM4Di-mCherry or AAV-DIO-mCherry (control) into the mPFC of 5xFAD mice. AAVretro-Cre and AAV-GRAB_gACh4m_ were also delivered to the DMS to achieve projection-specific expression of hM4Di and to enable ACh monitoring (Fig. 6A). Following expression of hM4Di, the 3-month-old mice received daily injections of the DREADD agonist, C21, six hours after instrumental training. This was repeated for four weeks, after which behavioral testing, electrophysiology recording, and Aβ immunostaining were performed.

**Figure 6.**
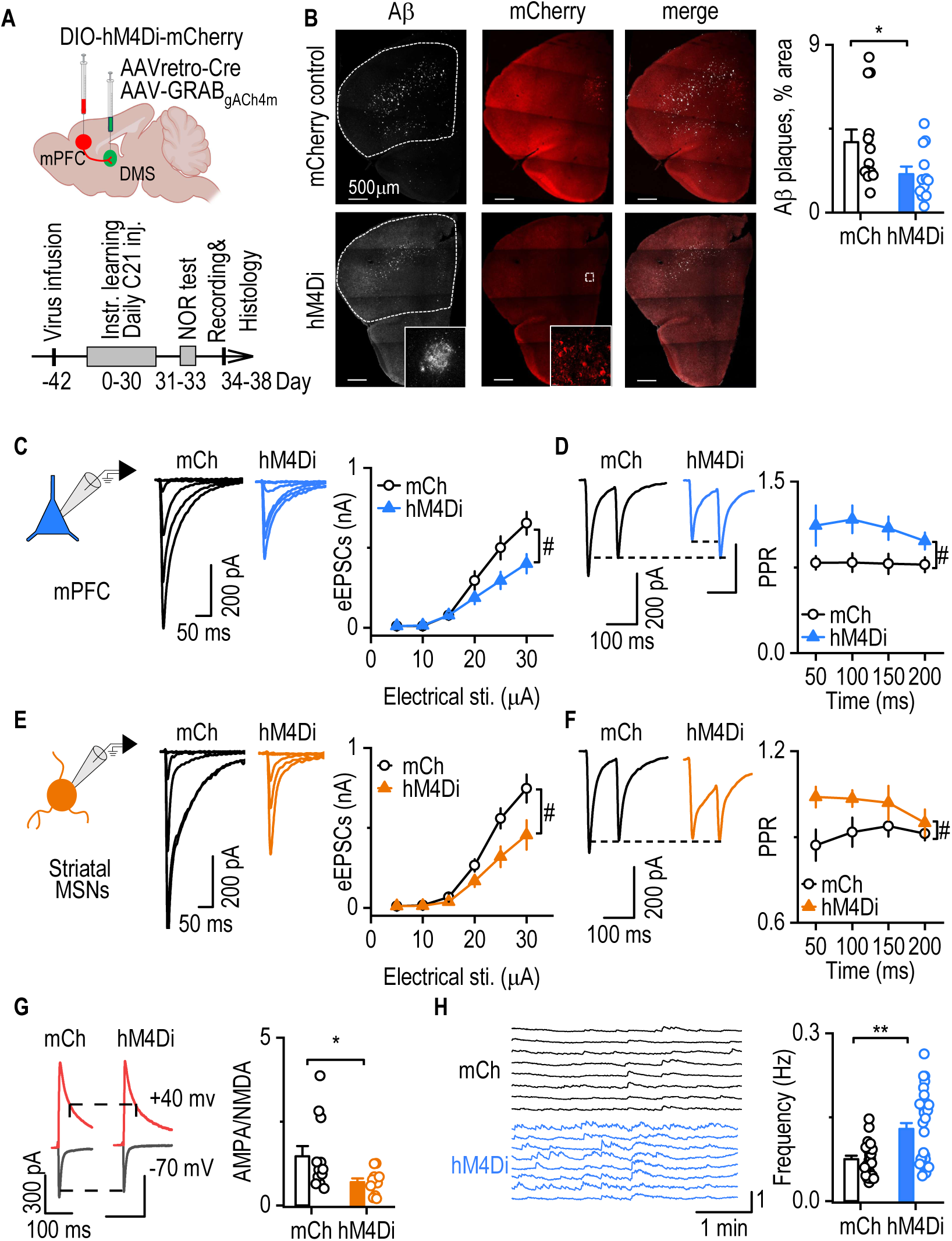
Sustained mPFC-to-DMS inhibition slows Aβ accumulation, reduces glutamatergic transmission, and enhances striatal ACh in 5xFAD mice. **A**, Schematic illustrating the viral infusion strategy and experimental timeline. At ∼2 months of age, 5xFAD mice were injected with either AAV-DIO-hM4Di-mCherry or AAV-DIO-mCherry in the mPFC, along with AAVretro-Cre and AAV-GRAB-gACh4m in the DMS. Mice received daily intraperitoneal injections of C21 (1 mg/kg) for four weeks after daily reversal learning training. Behavioral testing preceded electrophysiological recordings and Aβ immunostaining, which were conducted after the final C21 injection. **B,** Representative images of Aβ immunostaining in the mPFC and quantification region (dashed area). hM4Di-injected 5xFAD mice exhibited significantly reduced Aβ plaque coverage compared to mCherry-injected controls. Mann-Whitney Rank Sum Test, \**p* < 0.05. n = 13 mice (mCh) and 12 mice (hM4Di). **C,** eEPSCs amplitudes in mPFC-mCherry positive neurons were lower in the hM4Di group than in mCherry controls. Two-way RM ANOVA with Greenhouse–Geisser correction, ^#^*p* < 0.05. n = 17 neurons from 4 mice (mCh); 13 neurons from 4 mice (hM4Di). **D,** PPRs in mPFC-mCherry positive neurons were higher in hM4Di-injected mice than mCherry controls with Greenhouse–Geisser correction. Two-way RM ANOVA, ^#^*p* < 0.05. n = 15 neurons from 4 mice (mCh); 12 neurons from 4 mice (hM4Di). **E**, eEPSC amplitudes in DMS MSNs were smaller in hM4Di-injected 5xFAD mice than in mCherry controls. Two-way RM ANOVA with Greenhouse–Geisser correction, ^#^*p* < 0.05. n = 14 neurons from 4 mice (mCh); 13 neurons from 4 mice (hM4Di). **F**, PPR in DMS MSNs was higher in hM4Di-than in mCherry-injected 5xFAD mice. Two-way RM ANOVA with Greenhouse–Geisser correction, ^#^*p* < 0.05. n = 10 neurons from 4 mice (mCh); 8 neurons from 4 mice (hM4Di). **G,** AMPA/NMDA ratio was lower in hM4Di-than in mCherry-injected 5xFAD mice. Mann-Whitney Rank Sum Test, \**p* < 0.05. n = 14 neurons from 4 mice (mCh); 11 neurons from 4 mice (hM4Di). **H**, Sample traces of spontaneous striatal ACh release. The spontaneous ACh release frequency in DMS was higher in hM4Di-mice than in mCherry-injected 5xFAD mice. Mann-Whitney Rank Sum Test, \**p* < 0.05. n = 25 slices from 13 mice (mCh); 34 slices from 12 mice (hM4Di).

Cortical Aβ plaque analysis revealed a significantly lower plaque-covered area in hM4Di-injected mice, as compared to mCherry controls (Fig. 6B, *U* = 37, *p* = 0.028). This indicated that sustained inhibition of the mPFC-to-DMS circuit slowed Aβ accumulation. Electrophysiological recordings showed reduced excitatory drive in both the mPFC and DMS. Specifically, evoked EPSCs were smaller in mPFC neurons from hM4Di-injected mice (Fig. 6C, *F*_(1, 28)_ = 4.621, *p* = 0.0404), and PPRs were elevated (Fig. 6D, *F*_(1, 25)_ = 4.782, *p* = 0.0383), suggesting reduced presynaptic glutamate release in the mPFC. Similarly, DMS MSNs in hM4Di-injected mice exhibited lower evoked EPSC amplitudes (Fig. 6E, *F*_(1, 25)_ = 6.343, *p* = 0.0186), increased PPRs (Fig. 6F, F_(1, 16)_ = 4.695, *p* = 0.0457), and a lower AMPA/NMDA ratio (Fig. 6G, *U* = 31, *p* = 0.034). This indicated reduced excitatory transmission in the striatum.

To assess cholinergic output, we monitored spontaneous ACh release using the gACh4m sensor in the DMS. Sustained and selective inhibition of DMS-projecting mPFC neurons via hM4Di resulted in a significantly higher ACh event frequency compared to controls (Fig. 6H, U = 221, *p* = 0.0014). This indicated that sustained inhibition of DMS-projecting mPFC neurons increased striatal ACh levels.

To determine whether these effects extended beyond projection-specific inhibition to more widespread cortical suppression, we performed a parallel experiment using global mPFC inhibition by broadly expressing hM4Di in mPFC neurons (Supplementary Figure 8A). Global mPFC inhibition similarly reduced cortical Aβ accumulation (Supplementary Figure 8B-C), suppressed glutamatergic transmission in both the mPFC (Supplementary Figure 8D-J) and striatum (Supplementary Figure 9A-D), and increased striatal ACh release (Supplementary Figure 9E-L).

Taken together, our results indicate that mPFC-to-DMS hyperactivity drives cholinergic dysfunction in the striatum because sustained chemogenetic inhibition of the mPFC-to-DMS circuit slowed Aβ accumulation, reduced aberrant glutamatergic transmission, and restored striatal ACh levels.

### Sustained mPFC-to-DMS inhibition rescues reversal learning deficits in 5xFAD mice

Given our findings that mPFC-to-DMS circuit hyperactivity was associated with reduced striatal cholinergic tone, and previous data linking cholinergic deficits to impaired cognitive flexibility in aging^31^, we next examined whether sustained chemogenetic inhibition of this circuit could restore reversal learning deficits in 5xFAD mice.

We trained 5xFAD mice on an instrumental reversal learning task once daily for 30 days while maintaining inhibition of the corticostriatal circuit (Fig. 6A). To prevent neuronal inhibition from affecting memory consolidation, we administered C21 six hours post-training. During the initial learning phase, both hM4Di- and mCherry-injected 5xFAD mice exhibited similar lever-pressing rates (Fig. 7A, F_(1, 23)_ = 0.182, *p* = 0.675), and both groups preferentially pressed the valued over the devalued lever during the devaluation test (Fig. 7B, *Z* = -3.180, *p* < 0.001 for mCherry; *Z* = -3.509, *p* < 0.001 for hM4Di), confirming intact acquisition of the initial action-outcome contingency.

**Figure 7.**
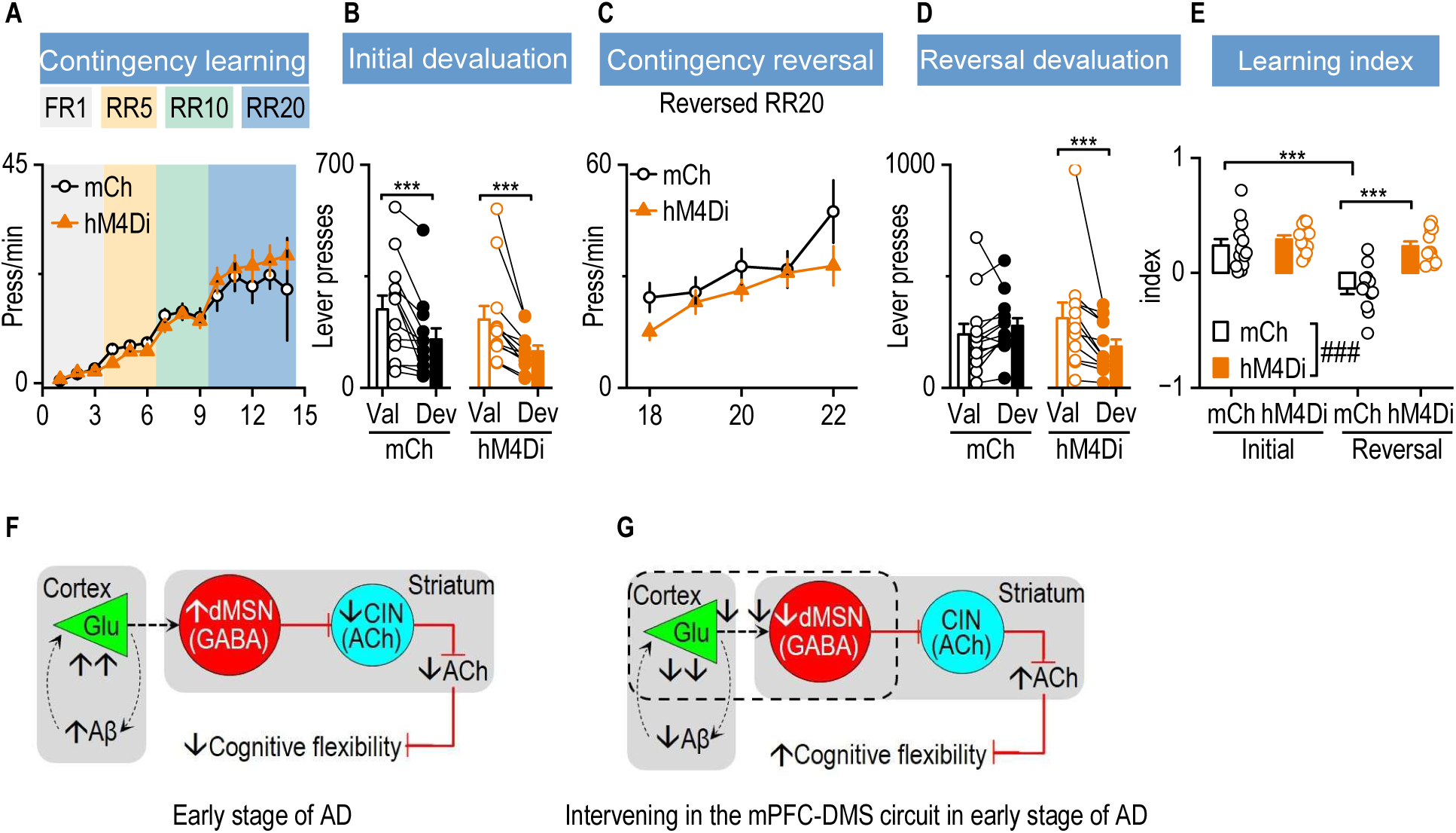
Sustained mPFC-to-DMS inhibition rescues reversal learning deficits in 5xFAD mice. **A**, hM4Di- and mCherry-injected 5xFAD mice exhibited similar lever-press rates during initial contingency learning, indicating comparable acquisition of the action-outcome association. Two-way RM ANOVA with Greenhouse–Geisser correction, ^#^*p* < 0.05. n = 13 (mCh) and 12 (hM4Di). **B,** Both mCh and hM4Di-injected mice displayed significantly fewer lever presses for devalued (Dev) than valued outcomes (Val) during the initial devaluation test. Wilcoxon Signed Rank Test, \*\*\**p* < 0.001. n = 13 (mCh) and 12 (hM4Di). **C,** During reversal contingency training, mCherry-(mCh) and hM4Di-injected (hM4Di) mice exhibited similar pressing rates. Two-way RM ANOVA with Greenhouse–Geisser correction. n = 13 (mCh) and 12 (hM4Di). **D**, hM4Di-injected (hM4Di) mice, but not mCherry-(mCh) 5xFAD, mice displayed significantly fewer lever presses for devalued than valued outcomes (Val) during the reversal devaluation test. Wilcoxon Signed Rank Test, \**p* < 0.05. n = 13 (mCh) and 12 (hM4Di). **E**, hM4Di-injected (hM4Di) mice exhibited a significantly higher reversal index than the reversal devaluation index of mCherry-injected (mCh) mice. The index is calculated as (Val-Dev)/(Val + Dev). Two-way RM ANOVA, ^###^*p* < 0.001; Sidak’s multiple comparisons post-hoc analysis, \*\*\**p* < 0.001. n = 13 (mCh) and 12 (hM4Di). **F and G**, Schematic models illustrating the proposed mechanism. In early-stage AD, hyperactivity in the corticostriatal circuit leads to impaired cognitive flexibility (F). Chronic inhibition of this circuit via chemogenetic inhibition restores cognitive flexibility in 5xFAD mice (G).

In the reversal phase, lever-pressing rates remained comparable between groups (Fig. 7C, *F*_(1,23)_ = 1.836, *p* = 0.189). However, hM4Di-injected mice showed significantly more presses for the valued lever in the reversal devaluation test, while mCherry-injected controls did not (Fig. 7D, *Z* = 1.922, *p* = 0.057 for mCherry; *Z* = -3.059, *p* < 0.001 for hM4Di). This indicated that only the treated group successfully acquired the reversed contingency. Importantly, hM4Di-injected mice exhibited a significantly higher reversal learning index than mCherry-injected controls (Fig. 7E; main effect: *F*_(1,23)_ = 37.974, *p* < 0.001), indicating improved cognitive flexibility following circuit inhibition.

*Post-hoc* comparisons further confirmed that the reversal learning index was significantly lower than the initial learning index in mCherry-injected mice (Fig. 7E, *t* = 4.529, *p* < 0.001), suggesting a reversal learning deficit in untreated 5xFAD mice. In contrast, hM4Di-injected mice showed no significant difference between their initial and reversal devaluation indices (Fig. 7E, *t* = 0.704, *p* = 0.488). This was consistent with the results observed in WT mice (Fig. 1G), indicating that long-term mPFC-to-DMS inhibition restored reversal learning performance in 5xFAD mice to near-WT levels. Critically, *post-hoc* comparisons also revealed that hM4Di-injected mice exhibited a significantly higher reversal learning index than the mCherry-injected controls (Fig. 7E, *t* = 5.352, *p* < 0.001). This suggested that sustained inhibition of the mPFC-to-DMS circuit effectively rescued the reversal learning deficits observed in young 5xFAD mice (Fig. 1 and Fig. 7).

We also evaluated broader cognitive performance using the novel object recognition test. While both groups preferred the novel object, hM4Di-injected 5xFAD mice displayed a significantly higher discrimination index than mCherry controls (Supplementary Figure 10), indicating selective inhibition of DMS projecting mPFC neurons enhanced recognition memory. Similarly, global mPFC inhibition by broadly expressing hM4Di in mPFC neurons also improved object recognition in 5xFAD mice (Supplementary Fig. 11A-D) without altering locomotor activity (Supplementary Figure 11E and F) in 5xFAD mice. These findings support the broader cognitive benefits of reducing mPFC hyperactivity.

Taken together, our data suggest that sustained mPFC-to-DMS inhibition rescues reversal learning deficits in 5xFAD mice.

## DISCUSSION

Cognitive flexibility is among the earliest executive functions impaired in AD. Here, we propose a circuit-based model in which early Aβ-induced cortical hyperactivity enhances mPFC-to-DMS excitatory drive, leading to hyperactivation of dMSNs, increased inhibitory control over CINs, reduced striatal ACh release, and impaired cognitive flexibility (Fig. 7F). Supporting this model, our findings demonstrate that sustained chemogenetic inhibition of the mPFC-to-DMS circuit during early-stage AD reduces cortical Aβ accumulation, normalizes glutamatergic transmission in both the mPFC and DMS, restores cholinergic tone, and ultimately rescues cognitive flexibility (Fig. 7G). These results highlight corticostriatal hyperactivity as an upstream driver of early cholinergic and cognitive deficits in AD and identify this circuit as a promising therapeutic target for early intervention.

Although striatal cholinergic dysfunction is linked to cognitive flexibility deficits in early-stage AD, the cholinergic hypothesis of AD focuses on the basal forebrain cholinergic system. This hypothesis is supported by postmortem studies showing significantly reduced cholinergic markers in the cortex and selective degeneration of the nucleus basalis of Meynert neurons in advanced AD^56, 57^. However, the striatum also contains large densities of CINs that modulate cognitive flexibility in goal-directed behaviors^22, 23, 25, 31^. Here, we report that 5xFAD mice exhibit deficits in striatum-dependent instrumental reversal learning, which precede the impairment of hippocampus-dependent spatial reversal learning. Our findings were consistent with a previous report that spatial learning deficits emerged at around six months of age in 5xFAD mice^36^. These observations suggest that deficits in cognitive flexibility are more sensitive to striatal dysfunction than to dysfunction in the basal forebrain cholinergic neurons, with the latter predominantly innervating the hippocampus. The critical role of striatal cholinergic activity in early-stage AD could relate to the high levels of AChE in the striatum^58^. Changes in striatal ACh, such as those caused by corticostriatal hyperactivity in early AD, likely impair striatum-dependent behaviors. These include executive functions such as instrumental reversal learning. The exact reasons why striatal, rather than basal forebrain, cholinergic activity is crucial in instrumental behavior inflexibility observed in early-stage AD remain unclear. However, our study suggested that CIN hypoactivity in 5xFAD mice was not due to cell loss, but resulted from excessive GABAergic inhibition by hyperactive dMSNs, which received elevated excitatory input from Aβ-enriched mPFC neurons. This multilevel circuit disinhibition cascade compromises striatal output and contributes to behavioral rigidity, evidenced by the early reversal learning deficits we observed in the 5xFAD mouse model.

Our study also raises intriguing questions about the interaction between CINs and MSNs under pathological conditions. Reduced ACh release may impair striatal neuron activity via direct and indirect pathways. Prior work suggested that ACh indirectly inhibited MSNs via GABAergic interneurons such as NPY-expressing neurons^27, 59, 60^, suggesting that CIN hypoactivity could lead to aberrant MSN recruitment during reversal learning. Additionally, elevated corticostriatal transmission may potentiate maladaptive memory trace reactivation, as enhanced input to engram neurons can artificially strengthen memory retrieval^61^. These mechanisms may converge to explain the excessive reactivation of cortical and striatal ensembles we observed during reversal learning in 5xFAD mice.

Mechanistically, our findings are consistent with the view that Aβ and neuronal hyperactivity form a pathological feedback loop. Activity-dependent Aβ release promotes further cortical hyperexcitability and impairs glutamate reuptake, thus enhancing glutamatergic transmission^10, 12, 53^. While Aβ accumulation is minimal in the striatum itself, the DMS receives dense glutamatergic input from Aβ-rich cortical areas. However, it is unclear how mPFC transmission to the striosome or matrix is altered under AD conditions. The elevated mPFC-to-dMSN transmission observed in the present study, therefore, impacts the striatum, despite the limited local deposition of Aβ. Our findings highlight the need to explore regions such as the striatum, which receives glutamatergic inputs from Aβ-rich cortical areas but is less affected by AD-related pathology.

Our study further demonstrates that sustained suppression of mPFC-to-DMS activity reduced cortical Aβ pathology. This was consistent with previous reports using hippocampal or entorhinal cortex regulation^54, 55, 62^. Moreover, we found that sustained mPFC-to-DMS suppression normalized excitatory transmission and resulted in elevated levels of ACh in the striatum. These effects were associated with improved reversal learning and recognition memory. Notably, general suppression of mPFC neurons produced a similar rescue of the circuit and behavior. This suggests that broad mPFC hyperactivity, rather than only within projection-defined subcircuits, can result in downstream dysfunction. Importantly, these effects persisted beyond the treatment period, indicating lasting circuit-level adaptations rather than transient pharmacological effects or simple learning-related changes. Early intervention to prevent Aβ accumulation in AD is important, as prolonged Aβ deposition can lead to lasting changes in corticostriatal circuits and even neuronal death, potentially limiting the clinical benefits of subsequent anti-Aβ treatments. While the development of anti-Aβ therapeutic antibodies continues, the findings of the present study indicate that reducing abnormal neuronal activity could provide a promising interim strategy for mitigating Aβ accumulation and reducing the associated cognitive deficits.

In summary, our study establishes a causal link between early Aβ-induced cortical hyperactivity, striatal cholinergic suppression, and cognitive inflexibility in the 5xFAD mouse model of AD neuropathology. Identifying the mPFC-to-DMS pathway as a modifiable driver of circuit and behavioral dysfunction provides a mechanistic framework for early executive decline in AD. These findings highlight the potential of circuit-based interventions to restore neuromodulatory balance and cognitive performance, particularly when applied at an early stage of disease progression before synaptic loss and neurodegeneration become widespread.

## MATERIALS AND METHODS

### Animals

ChAT-eGFP (stock # 007902), 5xFAD (stock # 034848), ArcTRAP (#021881), Ai14-tdTomato (#007914), and Drd1a-tdTomato (stock # 016204) mice were purchased from the Jackson Laboratory. D1-Cre mice were obtained from Mutant Mouse Regional Resource Centers. All mice were backcrossed onto a C57BL/6J background. ChAT-eGFP mice were crossed with 5xFAD to generate a 5xFAD;ChAT-eGFP line. 5xFAD;ChAT-eGFP mice were crossed with D1-Cre to generate D1-Cre;ChAT-eGFP;5xFAD line. D1-tdTomato mice were crossed with 5xFAD to generate a 5xFAD;D1tdT mouse line. ArcTRAP mice were crossed with Ai14 to generate Arc;Ai14 mice. Arc;Ai14 mice were crossed with 5xFAD mice to generate triple transgenic Arc;Ai14;5xFAD mice. Mice were group-housed at 23°C with a 12-h light:dark cycle (lights on at 11:00 pm). Food and water were provided ad libitum. Both male and female mice were used in this study. If there is no specific description, all mice for electrophysiology recordings were aged up to 4 or 5 months old. All animal care procedures and experimental protocols were approved by the Institutional Animal Care and Use Committee and were performed in agreement with the National Research Council Guide for the Care and Use of Laboratory Animals.

### Operant instrumental reversal learning tasks

This procedure was adapted from Bradfield and Balleine^31–33^, Ma and Huang^23^, and Gangal et al.^25^.

### Apparatus designs

The operant chambers (Med Associates Inc.) used in this study featured a central magazine for delivering rewards and two lever presses, one located on the left and the other on the right. A house light was also present. Each training session consisted of two sub-sessions, one involving the delivery of regular food pellets and the other involving purified food pellets, with the order of the sub-sessions selected randomly. A 2.5-minute time-out period separated the sub-sessions. Training sessions lasted a maximum of 62.5 minutes, with each sub-session ending at 30 minutes or when 20 rewards were earned, whichever came first. During the acquisition phase, the house light was on, and one of the lever presses was available for use, whereas during the time-out session, the house light was off, and all lever presses were retracted. The training schedule included four phases: initial learning, initial devaluation testing, reversal learning, and reversal devaluation testing.

### Acquisition of initial contingencies

Training began with either a purified or grain pellet sub-session, with the house light on and one of the lever presses presented. During initial training, the contingency is that the left lever (A1) results in a grain pellet reward (O1), and the right lever (A2) results in a purified pellet reward (O2). Animals were trained on a fixed ratio 1 (FR1, with one level press for one reward delivery) schedule for 3 days, where one lever press triggered one reward delivery. They then progressed to a variable ratio schedule, starting with a ratio of 5 (RR5, with a probability of 0.2 for reward delivery) for 3 days, followed by RR10 (with a probability of 0.1 for reward delivery) for 3 days, and finally reaching RR20 (with a probability of 0.05 for reward delivery) for five days.

#### Initial devaluation test

The devaluation test was conducted 24 hours after the last initial RR20 training. Before the test, animals were given one-hour free access to 40 randomly assigned purified or grain pellets to induce satiety. Ten minutes after the animal consumes all the pellets, or 60 minutes later, whichever comes first, the animals were placed into the operant chamber where the house light and both lever presses were presented. Lever presses during the 10-minute testing period were recorded and did not trigger reward delivery. The lever for the pre-feed pellet is considered a devaluated (Dev) lever, whereas the lever for a non-pre-feed pellet is considered a valued (Val) lever. The same devaluation test procedure for the other pellet type was conducted the following day.

#### Reversal learning

The animals underwent training on the reversal contingency after the initial devaluation test, beginning with a reversal-RR20 schedule. In this procedure, the action-outcome contingency was reversed from the initial action-outcome contingency, where the left lever (A1) was paired with purified pellet (O2), which was previously paired with grain pellet (O1), and the right lever (A2) was paired with grain pellet (O1), which was previously paired with purified pellet (O2), for five sessions.

#### Reversal devaluation test

The procedure for this devaluation test was the same as that for the initial devaluation test. The devaluation index is calculated as (Val-Dev)/(Val+Dev).

### ArcTRAP approach

The procedure was performed as previously described^63, 64^. The mice were habituated to receiving saline injections (i.p.) for three consecutive days before receiving 4-OHT (50 mg/kg, i.p.) injection. The 4-OHT solution was freshly prepared on the tagging day. In brief, the 4-OHT powder was dissolved in 200-proof ethanol (20 mg/mL) at room temperature, with continuous shaking until the 4-OHT was fully dissolved. An equal volume of a 1:4 mixture of castor oil and sunflower seed oil was added, resulting in a final concentration of 10 mg/mL 4-OHT, and the ethanol was removed by centrifugation under vacuum. The final 4-OHT in oil was stored at 4°C for a maximum of 12 h before use. The 4-OHT was administered immediately following either the completion of the middle two initial RR20 sessions or after the reversal RR20 sessions. Mice were first introduced to an isoflurane chamber immediately after the operant session, and 4-OHT was i.p. injected. The use of isoflurane is to reduce the pain and discomfort due to 4-OHT/oil mix injection. After the injection, mice were returned to their home-cage and left without any disturbance for the following 6 hours. Three weeks post-tagging, the mice were perfused. Subsequently, their brain slices were imaged using confocal microscopy.

### Confocal imaging and cell counting

Mice were intracardially perfused with 4% paraformaldehyde (PFA) in phosphate-buffered saline (PBS) 14 days after Arc;Ai14 labeling of activated neurons to allow sufficient time for replication and transcription^65^. The brains were extracted and post-fixed overnight in a 4% PFA/PBS solution, followed by dehydration in 30% sucrose. Each whole brain was sectioned serially into 50-µm coronal slices using a cryostat. Confocal images were obtained using a confocal laser-scanning microscope (Fluoview 3000, Olympus). Fluorescent images were reconstructed in three dimensions, and cell counts from these scans were manually acquired using Bitplane Imaris 8.3.1 (Bitplane, Zurich, Switzerland), as previously reported^66^. Neurons were counted using the Spot module within Imaris, which also calculated colocalization. To calculate the density of neurons, we conducted neuron counting in a circular region of interest (ROI) created in Imaris within the brain region. Neuron density is determined by dividing the number of neurons by the area of the defined ROI. All neuronal counts were performed with experimenters blinded to experimental conditions. Brain structures were registered using the Paxinos mouse atlas as a reference^67^.

### Barnes maze

The Barnes maze was performed as previously described^68^. The Barnes circular platform maze consists of a 66 cm diameter circular platform elevated on a 1.4 m stand. It features 20 evenly spaced 5.08 cm diameter holes around the circumference, with a black box (escape tunnel) positioned beneath one of the holes (San Diego Instruments, San Diego, CA, USA). Briefly, in each trial, the subject was initially placed at the center of the maze and given a maximum of 3 minutes to find the target escape hole. The trial ended when the mouse entered the escape hole. Spatial cues were present in the walls surrounding the maze to assist the animal in identifying the target location. A bright light was paired with the start of the experiment, and the light was turned off when the mouse reached the escape or at the end of each trial if the escape was not found. The test comprised four sessions: initial learning, probe testing, reversal learning, and reversal probe testing. During initial and reversal learning, mice underwent 4 consecutive training days, with 4 trials per day and a 15-minute intertrial interval. In the reversal learning session, the location of the target escape hole was shifted 180 degrees opposite of its position during initial learning. In both probe tests, the escape hole was removed, permitting mice to freely explore for 3 minutes. The entire test was monitored with an overhead camera and analyzed using video-tracking software (Ethovision XT, Noldus). The analysis included measures of velocity (cm/s), distance traveled (cm), immobility (%), and latency (s).

#### Locomotor activity test

The open-field test was conducted as previously described^69^. A transparent, open-field activity chamber (Med Associates, 43 cm x 43 cm x 21 cm height) was equipped with an infrared beam detector connected to a computer. Animals were moved to the locomotor activity test room 30 minutes before testing. The distance traveled and velocity were analyzed using Activity Monitor software (MED Associates, St Albans, VT). This test was conducted right after the last devaluation test of reversal instrumental learning, the final Barnes maze test, or 24 hours after the last C21 injection.

### Chemogenetic activation

A DREADD agonist, Compound 21 (C21) (dissolved in saline, 1 mg/kg, i.p.), was injected once every 24 hours for 30 continuous days at 3 months old of 5xFAD mice. For those 5xFAD animals that need to train with reversal learning, C21 injection was conducted 6 hours after instrumental training.

#### Novel Object Recognition Test

The test was comprised of three parts: habituation, familiarization, and testing.

##### Habituation

On the same day as the locomotor activity test, animals were given 10 minutes to explore an empty arena, an opaque, uncapped, cube-shaped chamber measuring 43 cm x 43 cm x 21 cm in height.

##### Familiarization

Twenty-four hours after habituation, the animals were exposed to the familiar arena with two identical objects for 10 minutes.

##### Testing

Twenty-four hours after familiarization, the mice were allowed to explore an open field containing both the familiar object and a novel object to assess long-term recognition memory. The test mouse was given 5 minutes to explore the chamber freely, and its behavior was monitored using an overhead camera and video-tracking software (Ethovision XT, Noldus). Ethovision was used to analyze the time spent in the ‘novel object zone’ and ‘familiar object zone,’ as well as the number of entries into each zone. The interaction zone is defined as the 2.5 cm area around each object.

#### Stereotaxic virus infusion

Stereotaxic viral infusions were performed as previously described^70–73^. The skin was opened to uncover the skull and expose the bregma and lambda, and the location of the desired injection site was determined. A three-axis micromanipulator was used to measure the spatial coordinates for bregma and lambda. Small drill holes were made in the skull at the appropriate coordinates, according to the Paxinos atlas. Brain infusions were performed at a rate of 0.1 µL/min. Two microinjectors were loaded with 0.5 µL of AAV2/9-hSyn-gACh4m (BrainVTA, PT-7021), AAV-hsyn-rACh1.7-WPRE-hGH-polyA (BrainVTA, PT-5488), or rAAV2Retro/CAG-Cre (UNC, AV7703H) were bilaterally injected into the striatum (AP: 0.26 mm, ML: ± 2.00 mm, DV: -3.50 mm). rAAV8/hSyn-hM4Di-mCherry (UNC, AV5360c), rAAV8/hSyn-mCherry (UNC, AV6443D), AAV9-hsyn-DIO-hM4Di-mCherry (Addgene, 44362), or AAV9-hsyn-DIO-mCherry (Addgene, 50459) were bilaterally injected into the mPFC region (AP: 1.94 mm, ML: ± 0.35 mm, DV: -2.20 mm) in 5xFAD or WT control mice. 0.5 µL of rAAV8/svn-ChR90-GFP(Svn-Chronos-GFP) (UNC, AV5842E) was infused bilaterally injected into the mPFC region in 5xFAD;D1-tdT or D1-tdT mice. 0.5 µL of rAAV9/syn-Flex-ChrimsonR-tdT (UNC, AV6556B) was infused bilaterally into the mPFC region in the DMS of D1-Cre;ChAT-eGFP; 5xFAD or D1-Cre;ChAT-eGFP mice. To avoid backflow of the virus, microinjectors were left in place for 10 min after the infusion was complete and then removed. The skin was sutured, and the mice were allowed to recover for at least 1 week prior to further experiments.

### Electrophysiological recordings of brain slices

#### Brain slice preparation

The brains were promptly removed, and slices of either the striatum or basal forebrain were cut 250 µm thick in an ice-cold cutting solution. The cutting solution consisted of the following (in mM): 40 NaCl, 143.5 sucrose, 4 KCl, 1.25 NaH_2_PO4, 26 NaHCO_3_, 0.5 CaCl_2_, 7 MgCl2, 10 glucose, 1 sodium ascorbate, and 3 sodium pyruvate, and had a pH of 7.35 and an osmolarity of 305-310 mOsm. The solution was saturated with 95% O_2_ + 5% CO_2_. Subsequently, the slices were incubated at 32°C for 45 minutes in a 1:1 mixture of the cutting and external solutions. The external solution, which had a pH of 7.35 and an osmolarity of 305-310 mOsm, was composed of the following (in mM): 125 NaCl, 4.5 KCl, 2 CaCl_2_, 1 MgCl_2_, 1.25 NaH_2_PO_4_, 25 NaHCO_3_, 15 sucrose, and 15 glucose. The external solution was also saturated with 95% O_2_ + 5% CO_2_. The slices were maintained in the external solution at room temperature until needed.

#### Electrophysiological recordings

The brain slice was positioned in a recording chamber attached to the fixed stage of an upright microscope (Olympus). The slice was then perfused with an oxygenated external solution at 32°C, with a 2 mL/min flow rate. Neurons were visualized using a 40x water-immersion lens and an infrared-sensitive CCD camera. A Multiclamp 700B amplifier with Clampex 10.6 software and Digidata1550A data acquisition system (Molecular Devices in Sunnyvale, CA) were employed for recording purposes. Recording pipettes with a resistance of 3-6 MΩ were made from a borosilicate glass capillary (World Precision Instruments, Sarasota, FL) by using a micropipette puller (Model P-97, Sutter Instrument Co).

To measure electrically evoked EPSCs in cortical neurons or dMSNs, a bipolar glass stimulating electrode was placed in the same compartment, and a brief pulse (40 mA) was delivered at 0.05 Hz. Cesium intracellular solution was used for EPSC recordings; this contained (in mM) 119 CsMeSO_4_, 8 TEA.Cl, 15 HEPES, 0.6 ethylene glycol tetraacetic acid (EGTA), 0.3 Na_3_GTP, 4MgATP, 5 QX-314.Cl and 7 phosphocreatine, with the pH adjusted to 7.3 using NaOH. For optically-evoked EPSC (oEPSC) recordings, 2 ms flash of light via objection lens was used (470 nm for Chronos and 590nm for Chrimson stimulation).

For the measurement of AMPAR/NMDAR ratio, the peak currents of AMPAR-mediated EPSCs were measured at a holding potential of –70 mV and the NMDAR-mediated EPSCs were estimated as the EPSCs at + 40 mV, 30 ms after the peak AMPAR EPSCs, when the contribution of the AMPAR component was minimal. The AMPA/NMDA ratio was calculated by dividing the AMPAR EPSCs by NMDAR EPSCs.

To measure electrically evoked IPSCs (eIPSCs) or optically evoked IPSCs (oIPSCs) in cholinergic interneurons (CINs), a high-chloride intracellular solution was used. This solution contained the following components (in mM): 125 CsCl, 6 NaCl, 10 HEPES, 1 EGTA, 10 QX-314.Cl, 2 MgATP, 6 Na₃GTP, and 2 Na₂CrPO₄. The pH was adjusted to 7.25, and the osmolarity was set at 280 mOsm. GABA currents (I_GABA_) in CINs were detected under whole-cell voltage-clamp mode (hold potential: -60 mV), and a high-chloride intracellular solution was used. To measure the I_GABA_ currents in CINs, CINs were held at -60mV in ACSF for 1 minute for baseline recording and 5 mL of GABA (50 μm) was applied. Holding currents were measured every 5s.

Cell-attached voltage-clamp or whole-cell current-clamp techniques were performed to record spontaneous action potentials of CINs without antagonists in the external solution. Whole-cell current-clamp was performed to record evoked firing. The pipette was filled with a K^+^-based intracellular solution consisting of 123 potassium gluconate, 10 HEPES, 0.2 EGTA, 8 NaCl, 2 MgATP, and 0.3 NaGTP (pH 7.25, 280 mOsm). Unlike whole-cell patch-clamp recordings, used to clamp neurons close to -70 mV, cell-attached patch-clamp was used to record neurons firing without a holding potential when the resistance exceeded 1 GΩ. Evoked action potentials were measured with whole-cell current-clamping while increasing current injections from 0 to 500 pA in incremental steps of 1-s.

Two continuous optical or electrical stimuli with fixed intervals (e.g., 50 or 100 ms), based on neuron type and experimental conditions, were applied to generate two post-synaptic currents. The peak amplitudes of the first (P1) and second (P2) responses were measured, and the paired-pulse ratio (PPR) was calculated as the ratio of P2 to P1. A minimum of three trials per condition were averaged for statistical analysis.

All patch-clamp recording data were analyzed using Clampfit (in pClamp 10.7, Molecular Devices), except that the sEPSCs/sIPSCs data were analyzed in the Mini Analysis Program.

#### Immunohistochemistry

Aβ immunostaining was conducted as previously described^74^. Antigen exposure was performed using 90% formic acid (Sigma, SHBQ5806). Non-specific proteins were blocked using 10% goat serum (Jackson ImmunoResearch, AB_2336990). Cortical sections of 50 μm thickness were labeled using rabbit anti-Aβ overnight at 4°C (Invitrogen, 71-5800, diluted at 1:300 in 2% goat serum of 0.5% PBS-Triton™ X-100) (Fisher BioReagents, BP151-500) (PBS-TX). This was followed by incubation with Alexa Fluor 647 conjugated goat anti-rabbit IgG (H + L) antibody (Invitrogen, A21245, diluted 1:500 in 2% goat serum of 0.5% PBS-TX) for 2 hours at room temperature and three washes in 0.5% PBS-TX. Quantifying the percentage area covered by Aβ plaques and the size of Aβ plaques was performed using ImageJ (NIH).

For the c-Fos immunostaining, sections were first placed into 0.5% PBS-TX (PBST) for 10 minutes, followed by blocking in 10% Bovine Serum Albumin serum (BSA) (Sigma, A7030) in PBS-TX for 1 hour and incubation in primary antibody (rabbit anti-cFos,1:2000, EMD-Millipore, cat# ABE457) diluted in blocking solution overnight under 4 degrees. Sections were then incubated in 1:500 biotinylated donkey anti-rabbit (Jackson Immuno Research #711-065-152) for 1 hour. Sections were visualized via incubation for 1 hour in Streptavidin-conjugated Alexa 647 (1:1000, Thermo Fisher, #21374, 2mg/ml).

#### Ex vivo live-tissue confocal imaging of ACh release

Acute brain slices were imaged in a custom-made chamber using an Olympus FluoView FV3000 confocal microscope, with ACSF flowing and saturated with 95% O_2_ and 5% CO_2_. The confocal microscope was equipped with a 10x NA 0.3 and a 40x NA 0.8 water immersion objective, along with a 488 nm and a 561 nm laser. The sample rate of imaging is 2-3 frames per second. Imaging parameters were consistently maintained across all imaging sessions, including laser intensity, HV, gain, offset, and aperture diameter. Electro-stimulation was administered using a glass pipette containing a tungsten filament. DF/F, z score, and half-width were analyzed and obtained by MATLAB. MATLAB scripts for data analysis are available upon request.

#### 6-Hz Seizure Threshold Testing

The 6-Hz model of partial seizures was employed to determine the seizure-inducing threshold in mice, following established protocols^75, 76^. Corneal stimulation consisted of monopolar rectangular pulses (0.2-millisecond duration) delivered at 6 Hz for 3 seconds using a constant-current device (World Precision Instruments, Sarasota, FL). To establish the EC50 value—the stimulation current required to induce seizures in 50% of animals—various current intensities ranging from 10 to 70 mA were administered to distinct groups of mice. A fixed current intensity of 50 mA was also used to allow direct comparison with the data obtained in the aforementioned studies (Supplementary Fig. 5). As an anti-pruritic, 0.5% tetracaine ocular anesthetic was applied to the corneas 10 minutes before stimulation. Just prior to stimulation, the corneal electrodes were moistened with 0.9% saline solution. Mice were manually restrained during the 6-Hz stimulation and released immediately afterward into an observation chamber. Behavioral seizure severity was assessed using the Racine scale^77^: stage 0: normal behavior; stage 1: chewing and facial twitches; stage 2: head nodding or shaking; stage 3: unilateral forelimb clonus without rearing, straub tail, and extended body posture; stage 4: bilateral forelimb clonus with rearing; stage 5: wild jumping and tonic-clonic activity. A minimum interval of 15 minutes was allowed before increasing the current intensity for subsequent stimulation. Seizures induced by the 6-Hz model typically lasted between 8 and 90 seconds, with mice resuming normal exploration within seconds of seizure cessation.

#### EEG electrode implantation

On the day of surgery, animals were weighed, anesthetized with a ketamine (100 mg/kg) and xylazine (10 mg/kg) mixture, and mounted on a stereotaxic frame. A vertical incision was made between the ears to expose the skull. Two wired anchor screws were implanted: one served as a surface electrode in the cortex and the other as a cerebellar reference electrode. The screws were secured to the skull with dental acrylic (P1 Technologies, Roanoke, VA). After a 10-day recovery period, the mice were single-housed in the vivarium prior to the pilocarpine challenge. Cortical recordings were made from the animals.

#### Pilocarpine seizure test

Animals were pretreated with scopolamine methyl bromide (1 mg/kg, subcutaneous, Sigma-Aldrich, St. Louis, MO) 30 minutes prior to pilocarpine injection to enhance survival rates without affecting seizure severity. Pilocarpine (416 mg/kg, intraperitoneal, Sigma-Aldrich, St. Louis, MO) was then administered to induce persistent seizures and status epilepticus in the mice^78^.

#### Video EEG and Behavior Recording

Behavior and EEG activity were continuously monitored for 2 hours after pilocarpine injection to assess seizure progression. A 30-minute baseline EEG recording was conducted prior to the experiment using a digital EEG system. EEG signals were recorded with ClampEx-pCLAMP software connected to Grass Technologies P511 AC preamplifiers, sampled at 2000 Hz, and filtered through a Digidata 1440A to reduce noise. Video recordings of the EEG analysis were made with 1080p HD digital security cameras equipped with infrared LEDs for night vision. Both cortical and hippocampal EEG channels were recorded. Mice were awake and freely moving in cages fitted with swivels. Pilocarpine-induced EEG seizure activity was characterized by high-amplitude discharges (at least twice the baseline amplitude) and repetitive discharges (>0.5 Hz). The severity of behavioral seizures was rated using the Racine scale described earlier.

### Statistical analysis

Data were analyzed by a two-tailed t-test or 2-way RM ANOVA followed by Sidak post-hoc analysis. Normality and equal variance assumptions were tested. Mann-Whitney U test, Wilcoxon Signed-Rank test, unpaired t-test with Welch’s correction, mixed-effects models or Generalized Linear Mixed Models (GLMM) were used when normality or equal variance assumptions was violated. Greenhouse–Geisser correction was applied when the sphericity assumption may not hold in the 2-way RM ANOVA. Significance was determined if *p* < 0.05. Statistical analysis was conducted by SigmaPlot, Prism or SPSS. Data are presented as mean ± s.e.m.

## Acknowledgments

This research was supported by TRSA John P. McGovern Fellowship 2024 (Y.H) and by NIH grants U01AA025932 (J.W.), R01AA027768 (J.W.), and R01AA030293 (J.W.)

## Author contributions

J.W. and Y.H. conceived the project and designed the experiments. Y.H., X.X. and K.S. performed the behavioral experiments and analyzed the corresponding data. Y. H., Z.H., and H.G. performed the electrophysiological experiments and analyzed the corresponding data. Y.H., X.X., J.H., and X.W. performed the histology experiments. Y.H. and R.C. performed the live imaging confocal experiments and analyzed the corresponding data. X. W. and D.S.R performed the seizure experiments and analyzed the data. X.W. performed animal breeding for all experiments. J.W., and Y.H. wrote the manuscript with substantial input from J.C.

## Supplementary Figures

**Supplementary Figure 1.**
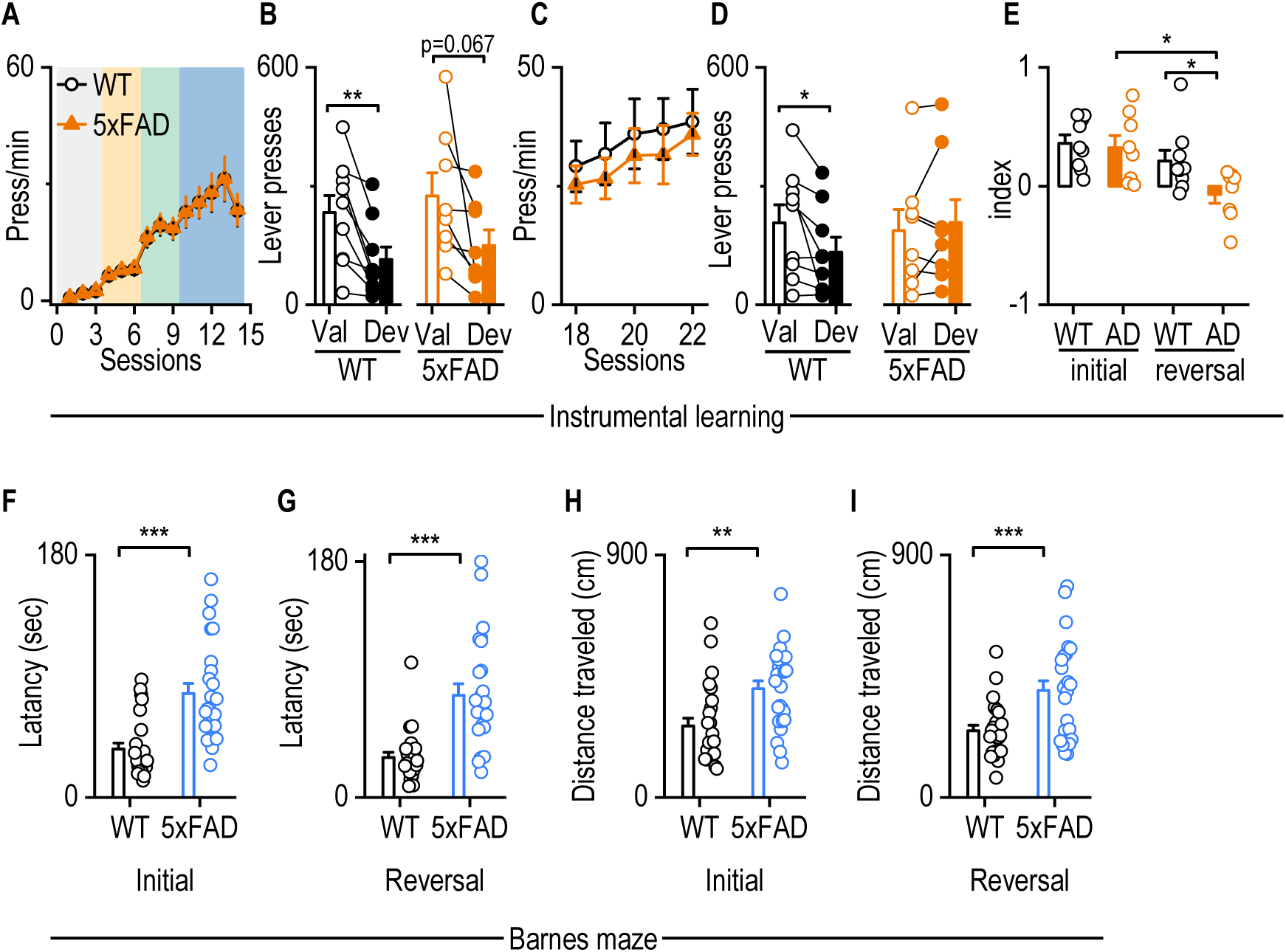
5xFAD mice exhibited reversal learning deficits in instrumental learning and Barnes maze test at 6 months of age. **A,** 6-month-old WT and 5xFAD mice did not differ in lever pressing rates during the initial contingency training. Two-way RM ANOVA with Geisser-Greenhouse correction. F_(1,15)_ = 0.266, *p* = 0.614. n = 9 (WT); 8 (5xFAD). **B,** WT mice displayed fewer lever presses for devalued than valued outcomes during the initial devaluation test, and 5xFAD mice exhibited a similar trend. Paired t-test, t_8_ = 4.059, p = 0.0036 for WT; t_8_ = 2.169, p = 0.067 for AD. \*\**p* < 0.01. n = 9 (WT); 8 (5xFAD). **C,** WT and 5xFAD animals had similar lever pressing rates during reversed contingency training. Two-way RM ANOVA with Geisser-Greenhouse correction, F_(1,15)_ *=*0.305*, p* = 0.589. n = 9 (WT); 8 (5xFAD). **D**, WT, but not 5xFAD, mice displayed fewer lever presses for devalued than valued outcomes during the reversal devaluation test. Paired t-test, t_8_ = 2.569, p = 0.0332 for WT; t_7_ = 0.843, p = 0.427 for AD. \**p* < 0.05. n = 9 (WT); 8 (5xFAD). **E,** 5xFAD mice displayed a significantly lower reversal index than their initial devaluation index and the reversal devaluation index of WT mice. Two-way RM ANOVA with Geisser-Greenhouse correction, F_(1,15)_ = 4.504, p = 0.051; Sidak’s multiple comparisons post-hoc analysis, *p < 0.05 for sessions within AD. \**p* < 0.05. n = 9 (WT); 8 (5xFAD). **F and G,** 6-month-old 5xFAD mice spent more time finding the escape box than WT mice did in both initial (F, U = 90, *p* < 0.001) and reversal sessions (G, U = 78, *p* < 0.001). Mann Whitney test, \*\*\**p* < 0.001. n =25 WT and 5xFAD. **H and I,** The total traveled distance from the starting point to the escape box by 6-month-old 5xFAD mice was higher than that of WT mice in both initial (H, *U* = 144, *p* < 0.01) and reversal sessions (I, *t*_48_ = -3.538, *p* < 0.001). Mann Whitney test or unpaired t-test, ** *p* < 0.01, \*\*\**p* < 0.001. n =25 WT and 5xFAD.

**Supplementary Figure 2.**
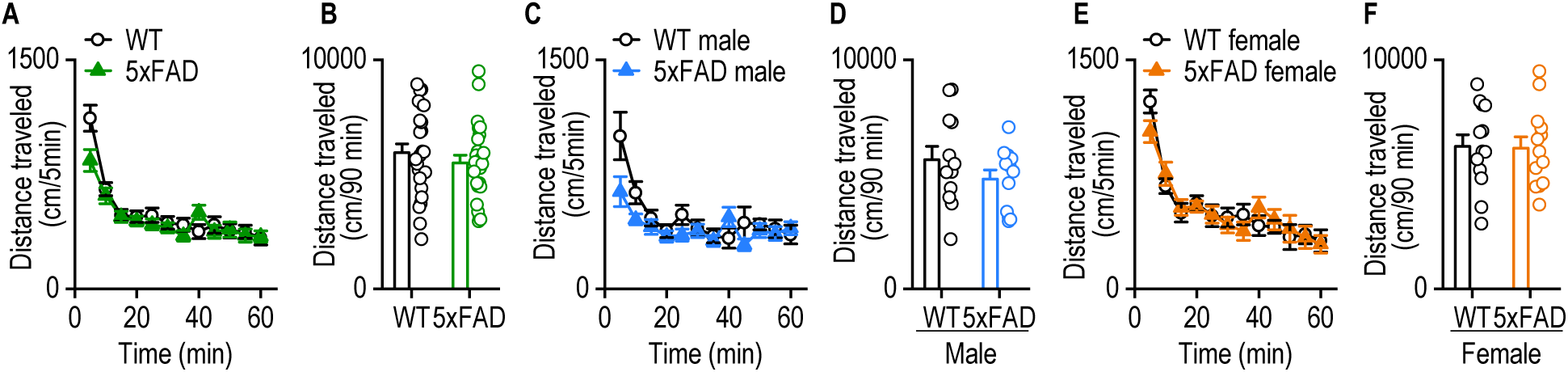
5xFAD mice did not exhibit abnormal locomotor activity. **A**, **B**, Similar spontaneous locomotor activities were shown in WT and 5xFAD mice in time-course (A, *F*_(1, 48)_ = 0.7615, *p* = 0.3872) or the total traveled distance (B, *t*_48_ = 0.873, *p* > 0.05). Two-way RM ANOVA and unpaired t-test. n = 25 WT and 5xFAD. **C-F**, There were no differences in spontaneous locomotor activities between males (C, *F*_(1, 22)_ = 1.466, *p* = 0.2388; D, *t*_22_ = 1.211, *p* > 0.05) or females (E, *F*_(1, 24)_ = 0.009951, *p* = 0.9214; F, *t*_24_ = 0.0988, *p* > 0.05) 5xFAD mice and their WT control. Two-way RM ANOVA and unpaired t-test. n = 12 male WT and AD; 13 female WT and 5xFAD.

**Supplementary Figure 3.**
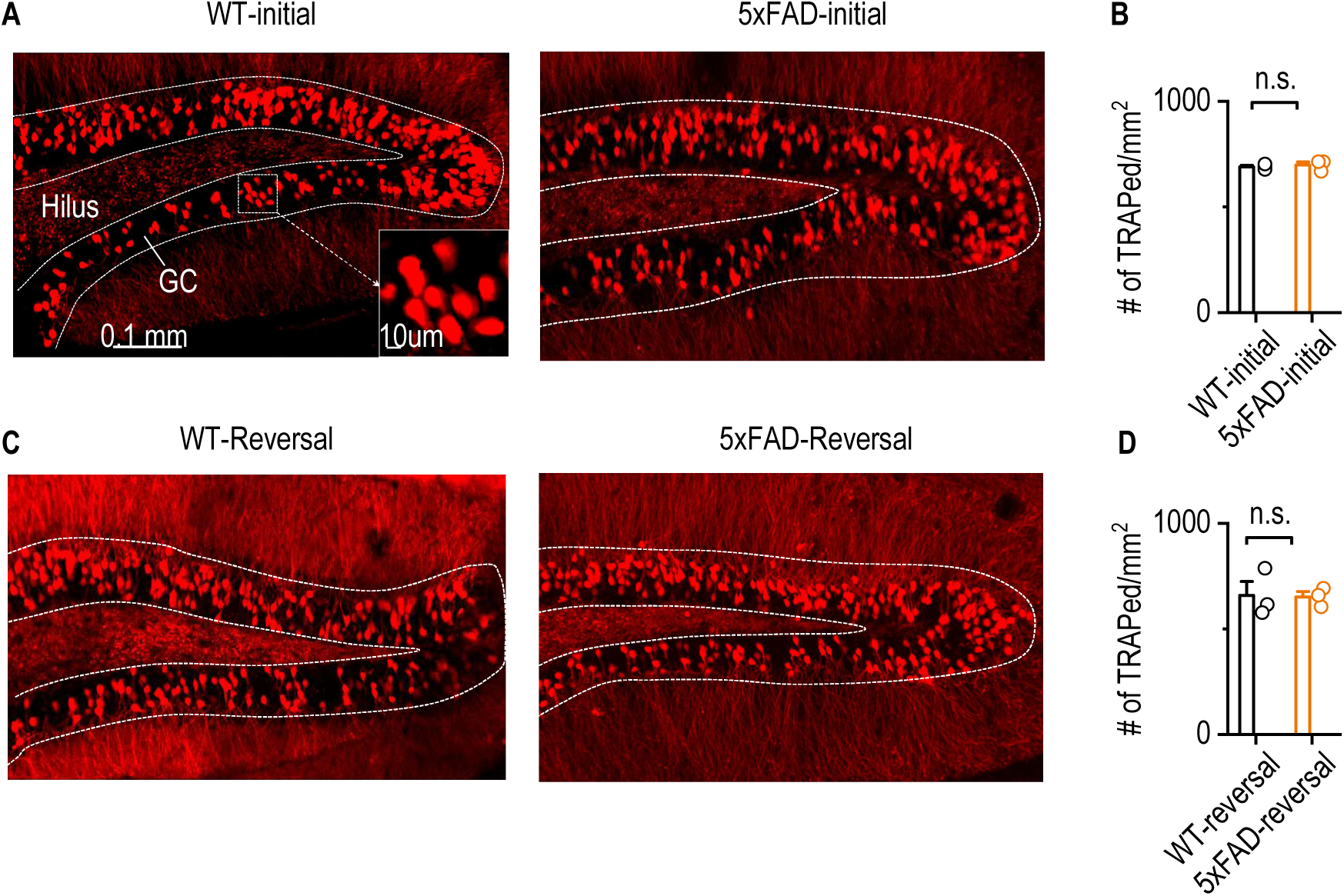
Trapped hippocampus neurons do not differ between WT and 5xFAD in either initial or reversal learning. **A,** Representative images showing trapped neurons in the dentate gyrus (DG) of the hippocampus following initial learning in WT and 5xFAD mice. GC, granule cell layer. **B,** Quantification of tdTomato-labeled DG neurons during initial learning revealed no significant difference between WT and 5xFADmice. t_4_ = 0.5449, *p* = 0.6148, unpaired t-test. n = 3 mice for both WT and 5xFAD. **C,** Representative images showing trapped DG neurons during reversal learning in WT and 5xFAD mice. **D,** Quantification of labeled DG neurons during reversal learning showed no significant difference between WT and 5xFAD mice. t_4_ = 0.1146, *p* = 0.9143, unpaired t-test. n = 3 mice for both WT and 5xFAD.

**Supplementary Figure 4.**
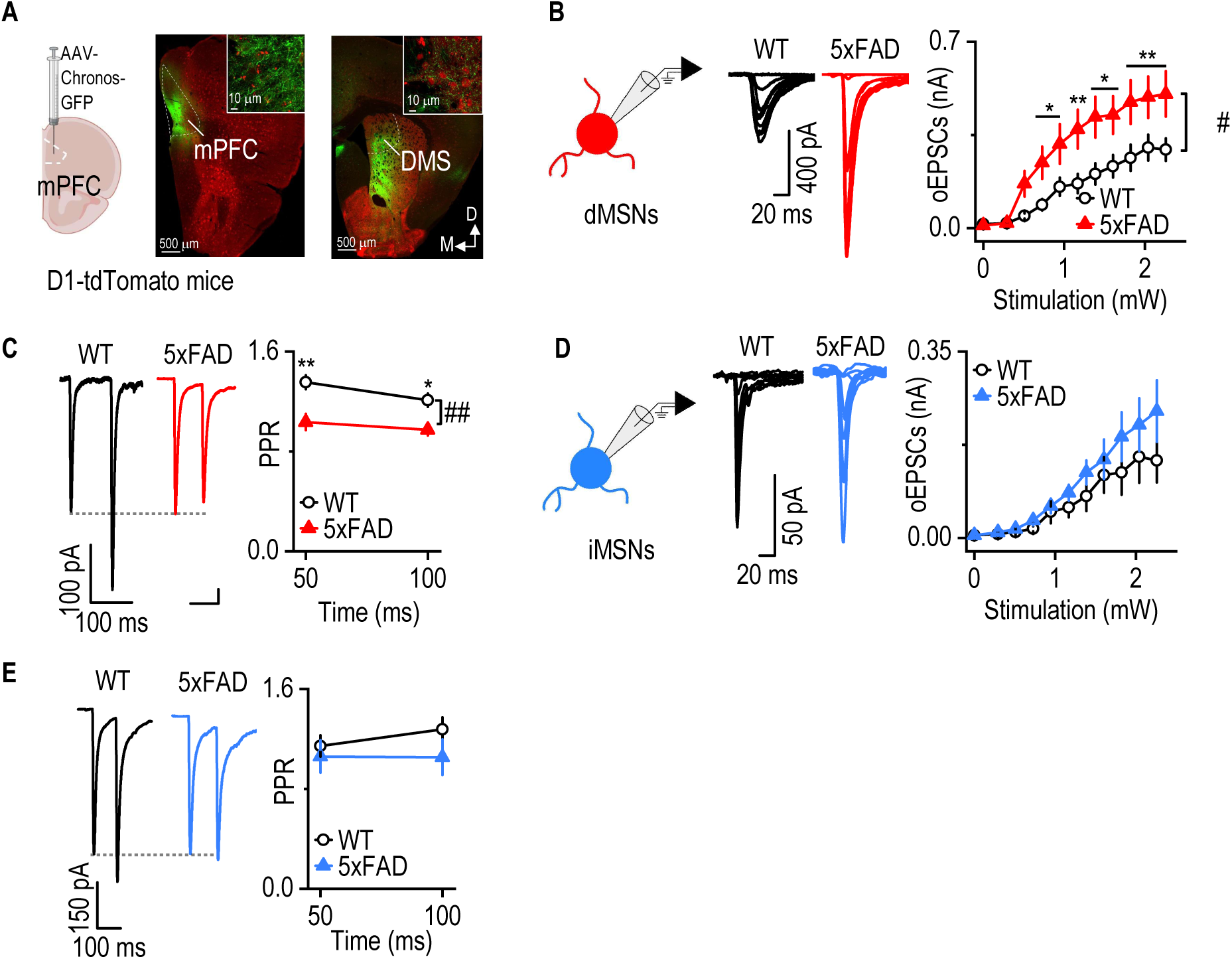
The mPFC-to-dMSN circuit is hyperactive in 12-month-old 5xFAD mice. **A,** Representative image showing Chronos-eGFP expression in the injection site (mPFC) and its projection to the DMS in 5xFAD;D1-tdTomato mice. **B,** oEPSC amplitude in DMS dMSNs was greater in 12-month-old 5xFAD mice than WT controls. Two-way RM ANOVA, *F*_(1,32)_ = 5.422, *p* = 0.026, ^#^*p* < 0.05; Sidak’s multiple comparisons post-hoc analysis, \**p* < 0.05, \*\**p* < 0.01, versus WT at the same stimulating intensities. N = 17 neurons from 3 mice (WT and 5xFAD). **C**, PPRs oEPSCs in DMS dMSNs were smaller in 12-month-old 5xFAD mice than age-matched WT controls. Two-way RM ANOVA, *F*_(1,27)_ = 9.080, *p* = 0.006, ^##^*p* < 0.01; Sidak’s multiple comparisons post-hoc analysis, \*\**p* < 0.01, versus WT at the same time interval. n = 15 neurons from 3 mice (WT) and 14 neurons from 3 mice (5xFAD). **D,** oEPSC amplitudes in DMS iMSNs were not significantly different between 12-month-old 5xFAD and WT mice. Two-way RM ANOVA, *F*_(1, 19)_ = 1.167, *p* = 0.2935. n = 9 neurons from 2 mice (WT) and 12 neurons from 3 mice (5xFAD). **E**, PPRs of oEPSCs in iMSNs did not differ between 5xFAD and WT mice. Two-way RM ANOVA, *F* _(1, 14)_ = 0.07133, *p* = 0.7933. N = 6 neurons from 2 mice (WT) and 10 neurons from 3 mice (5xFAD).

**Supplementary Figure 5.**
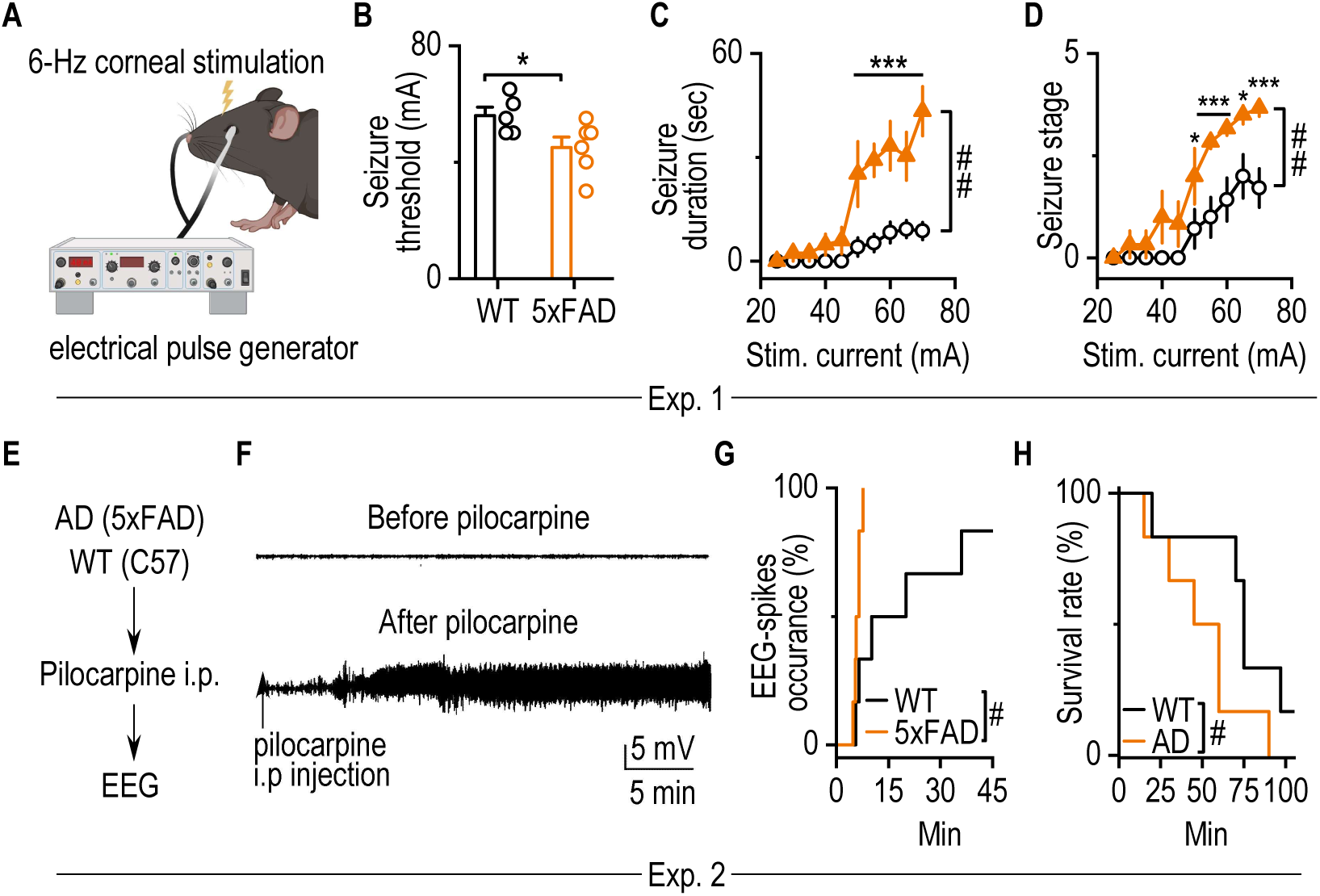
5xFAD mice exhibit heightened induced seizure activity. A, Schematic illustrating 6-Hz corneal stimulation in mice. Corneal stimulation consisted of monopolar rectangular pulses (0.2-millisecond duration) delivered at 6 Hz for 3 sec using a constant-current device. **B,** 5xFAD mice displayed a significantly lower threshold for seizure induction compared to WT controls. \**p* < 0.05, unpaired t-test. n = 6 (5xFAD) and 7 (WT) mice. **C,** 5xFAD mice had significantly longer seizure durations than WT controls. ^##^*p* < 0.01; \*\*\**p* < 0.001 versus WT at the same stimulating intensities, two-way RM ANOVA. n = 6 (5xFAD) and 7 (WT) mice. **D,** 5xFAD mice exhibited increased seizure stages compared to WT controls. ^##^*p* < 0.01; \**p* < 0.05, \*\*\**p* < 0.001 versus WT at the same stimulating intensities two-way RM ANOVA. n = 6 (5xFAD) and 7 (WT) mice. **E,** Schematic of the experimental design for an additional cohort of AD and WT mice in Exp. 3. Pilocarpine was administered intraperitoneally to induce seizure behavior, and EEG spikes were recorded from the cortex. **F,** Sample trace in EEG recordings showing spikes before and after pilocarpine i.p. injection. **G,** 5xFAD mice displayed a higher percentage of seizure spike occurrences in EEG recordings within 45 min after pilocarpine injection compared to WT controls. ^#^*p* < 0.05, Kaplan-Meier estimates followed by the Tarone-Ware test. n = 6 mice per group. **H,** Survival rate following seizure induction was significantly lower in 5xFAD mice compared to WT controls. ^#^*p* < 0.05, Kaplan-Meier estimates followed by the Tarone-Ware test. n = 6 mice/group.

**Supplementary Figure 6.**
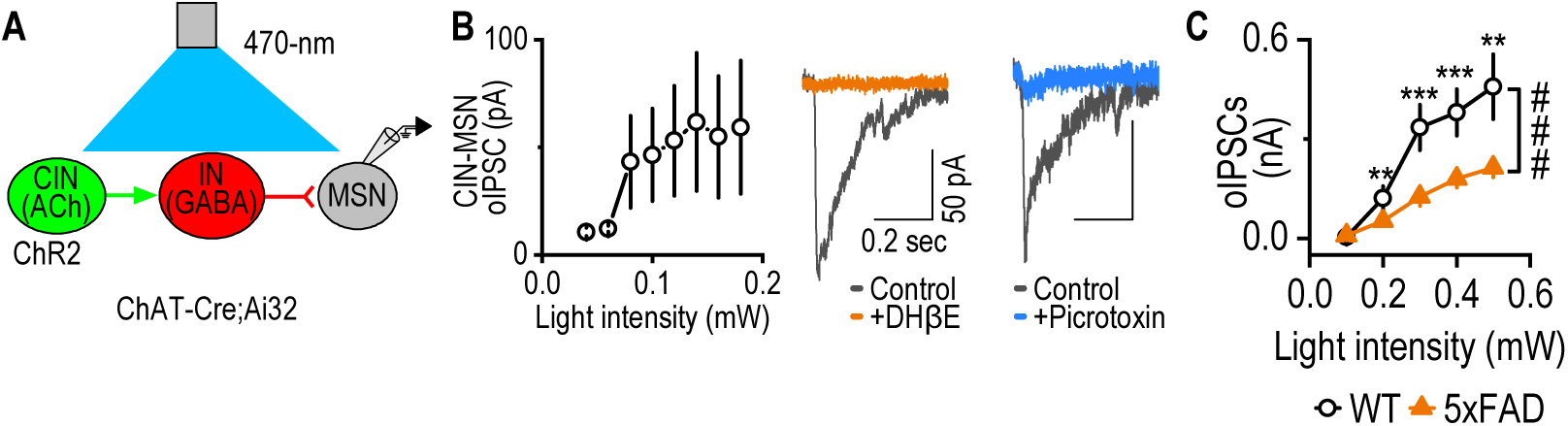
Reduced CIN-to-MSN inhibition in 5xFAD mice. **A,** Schematic illustrating the stimulation and recording of CIN-to-MSN inhibitory transmission in DMS slices from 4-month-old ChAT-Cre;Ai32 mice. IN, interneuron. **B,** Representative CIN-to-MSN inhibitory postsynaptic currents (IPSCs, left) were blocked by the nicotinic receptor antagonist DhβE (1 µM) and the GABA_A_ receptor antagonist picrotoxin (0.1 mM), confirming both cholinergic and GABAergic components. n = 7 neurons from 2 mice. **C,** CIN-to-MSN IPSC amplitudes were significantly reduced in 5xFAD mice compared to controls. *F*_(1, 39)_ = 24.72, ^###^*p* < 0.001, Two-way RM ANOVA; \*\**p* < 0.01, \*\*\**p* < 0.001 versus AD at the same stimulating intensities, Sidak’s multiple comparisons post-hoc analysis. n = 20 neurons from 3 mice (Ctrl; ChAT-Cre;Ai32) and 21 neurons from 3 mice (5xFAD;ChAT-Cre;Ai32;5xFAD).

**Supplementary Figure 7.**
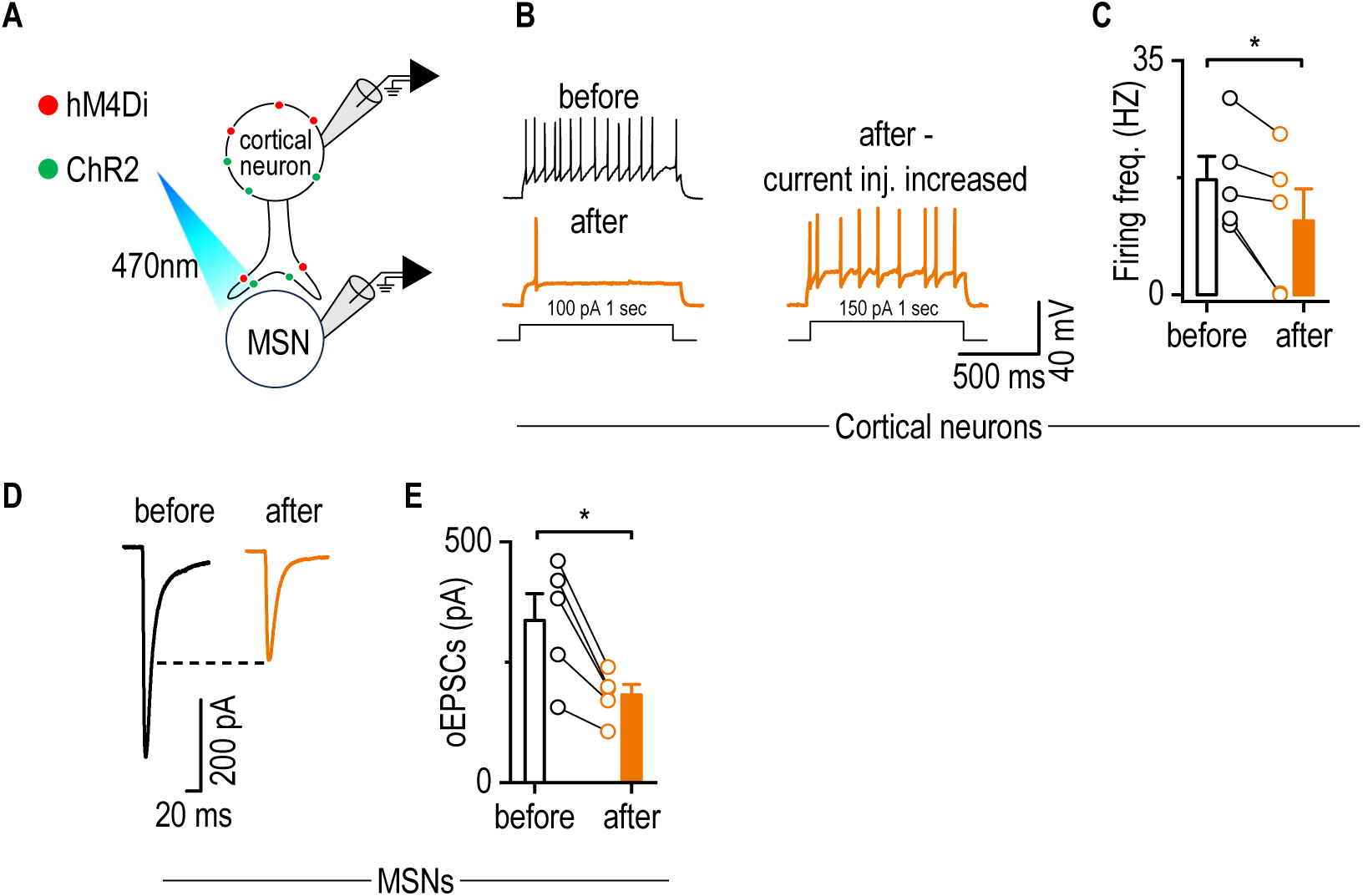
hM4Di activation reduces neuronal excitability and synaptic transmission. **A,** Schematic illustrating AAV infusion and patch-clamp recording strategy. AAV-hSyn-ChR2-GFP and AAV-hM4Di-mCherry were co-infused into the mPFC. Recordings were obtained from mPFC neurons co-expressing ChR2 and hM4Di, and from postsynaptic MSNs in the DMS that did not express either construct. **B,** Sample trace showing decreased excitability of hM4Di-expressed mPFC neuron after hM4Di activation. **C,** hM4Di activation significantly reduced excitability. Paired t-test, \**p* < 0.05. **D,** Sample trace showing decreased oEPSC amplitude after hM4Di activation. **E,** hM4Di activation significantly reduced oEPSC amplitude. Paired t-test, \**p* < 0.05.

**Supplementary Figure 8.**
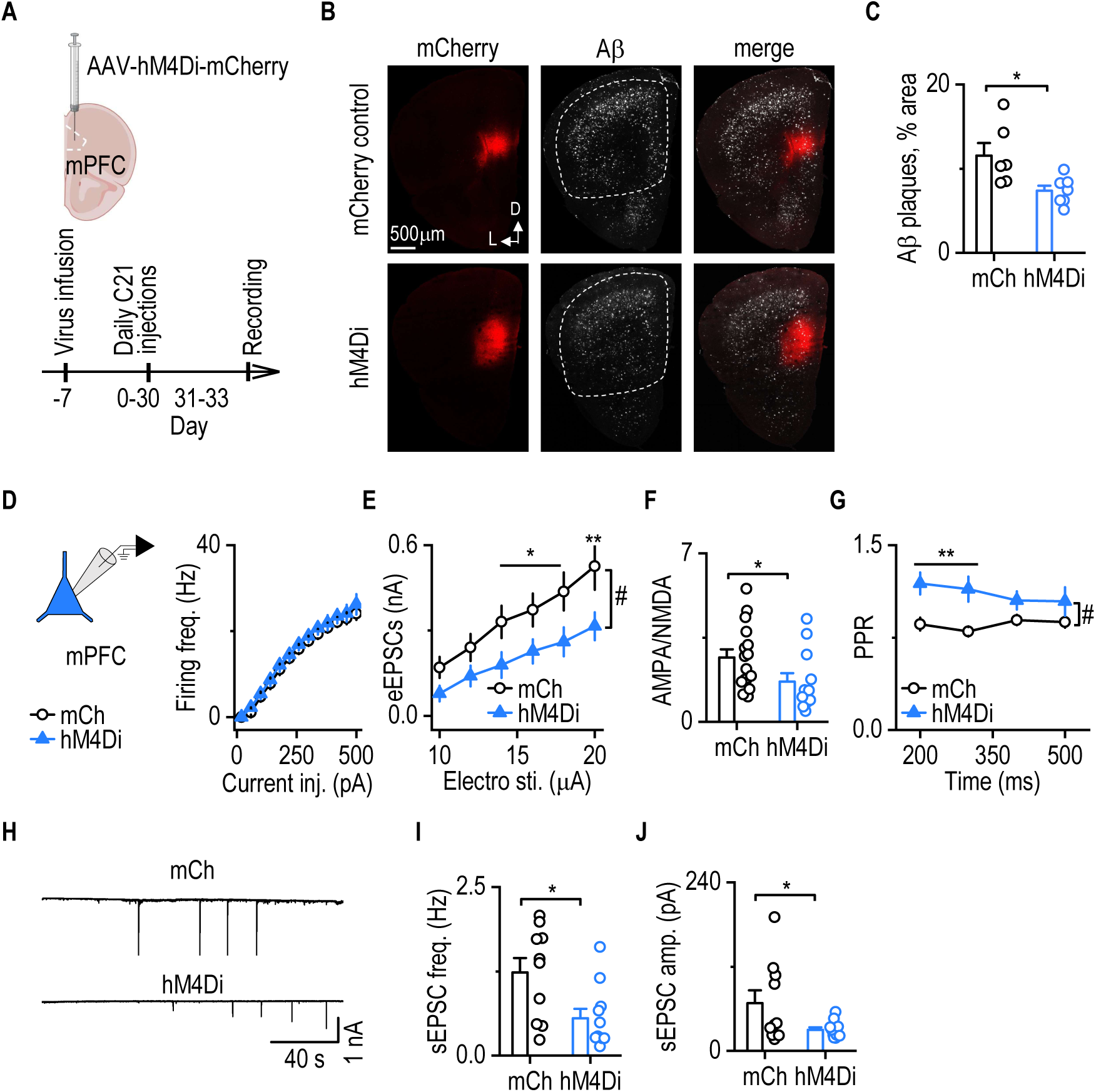
Sustained chemogenetic inhibition of cortical neurons normalizes glutamatergic transmission and results in lower Aβ accumulation in the cortex. **A**, Schematic illustrating the experimental timeline. At 3 months of age, 5xFAD mice were infused with either AAV-hM4Di-mCherry or AAV-mCherry in the mPFC and received daily intraperitoneal C21 injections (1 mg/kg) for four weeks. Behavioral testing was conducted before brain slices were collected for electrophysiological recordings and Aβ staining following the final C21 injection. **B,** Representative images showing Aβ staining and quantification regions (dashed area) in hM4Di- and mCherry-injected 5xFAD mice. **C,** The percentage of area covered by Aβ plaques was significantly lower in hM4Di-injected mice than in mCherry-injected controls. unpaired t-test, *t*_11_ = 2.751, *p* = 0.0188. \**p* < 0.05. n = 6 mice (mCh); 7 mice (hM4Di). **D,** Excitability of mCherry-positive mPFC neurons did not differ between hM4Di and mCherry (mCh) groups. Two-way RM ANOVA, *F*_(1,24)_ = 0.068, *p* = 0.797. n= 13 neurons from 4 mice (mCh and hM4Di). **E,** eEPSC amplitudes in mCherry-positive mPFC neurons were significantly lower in hM4Di-injected mice than in mCherry controls. Two-way RM ANOVA, F_(1,32)_ = 4.835, *p* = 0.035, ^#^*p* < 0.05; Sidak’s multiple comparisons post-hoc analysis, \**p* < 0.05, \*\**p* < 0.01 versus mCh at the same stimulating intensities. n = 15 neurons from 4 mice (mCh) and 19 neurons from 4 mice (hM4Di). **F,** AMPA/NMDA ratios were lower in hM4Di-injected mice than in mCherry controls. Unpaired t-test. *t*_26_ = 2.048, *p* = 0.051. \**p* < 0.05. n = 16 neurons from 4 mice (mCh) and 12 neurons from 4 mice (hM4Di). **G,** PPRs of mCherry-positive mPFC neurons were higher in hM4Di-injected mice than in mCherry controls. Two-way RM ANOVA, *F*_(1,33)_ = 7.153, *p* < 0.05; ^#^*p* < 0.05; Sidak’s multiple comparisons post-hoc analysis, \*\**p* < 0.01 versus WT at the same time interval. n= 16 neurons from 4 mice (mCh) and 19 neurons from 4 mice (hM4Di). **H,** Representative traces of sEPSCs recorded from mCherry-positive mPFC neurons in both groups. **I and J,** Both the frequency and amplitude of sEPSCs in mPFC-mCherry positive neurons were lower for hM4Di-than mCherry-injected 5xFAD mice. unpaired t-test, *t*_19_ = 2.691, *p* = 0.0145(**G**);*t*_19_ = 2.170, *p* = 0.0429; \**p* < 0.05. n= 10 neurons from 4 mice (mCh) and 11 neurons from 4 mice (hM4Di).

**Supplementary Figure 9.**
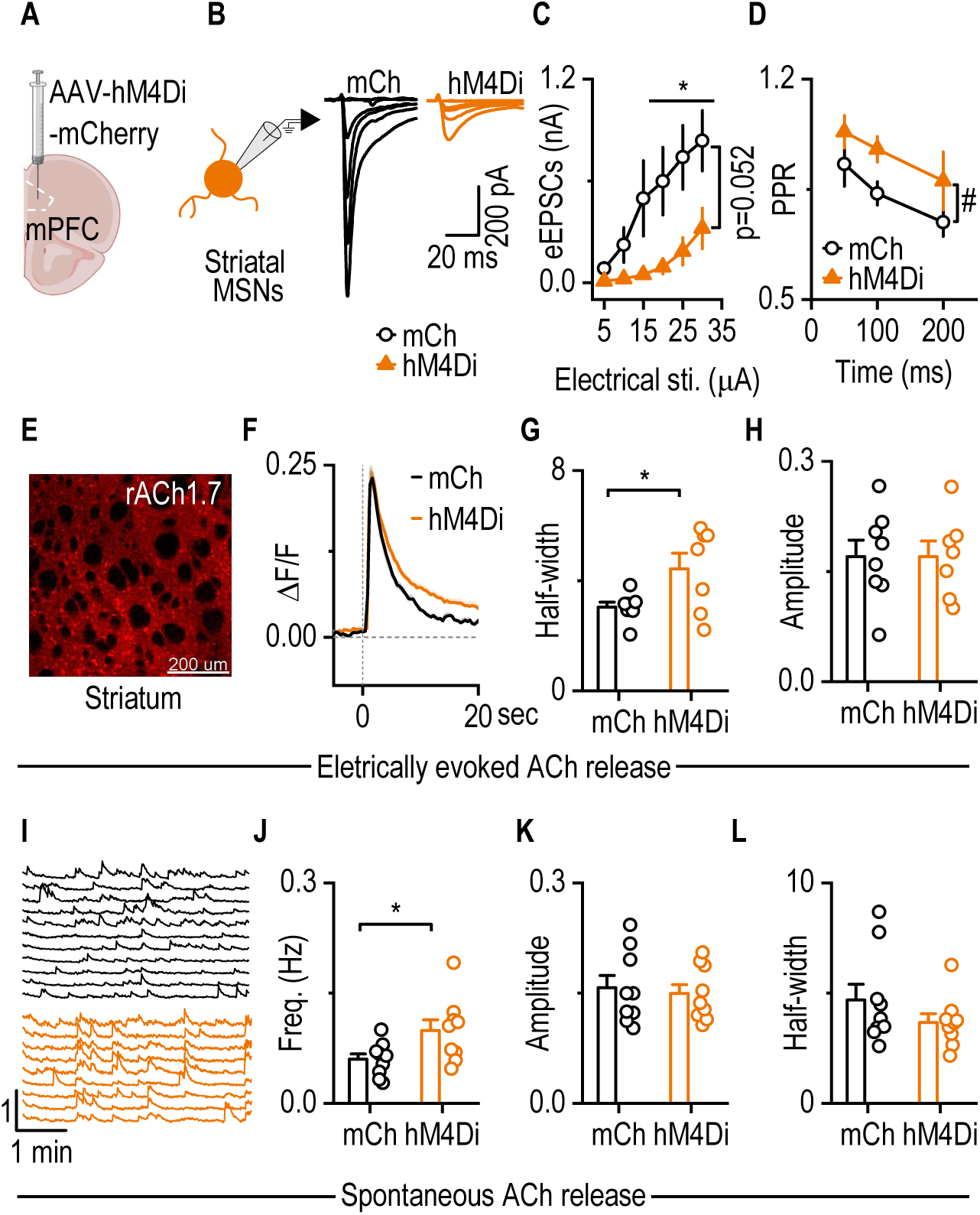
Sustained chemogenetic inhibition of cortical neurons leads to decreased glutamatergic transmission and increased striatal ACh levels. **A**, Schematic showing AAV-hM4Di-mCherry or AAV-mCherry infusion into the mPFC. **B,** Representative trace of eEPSCs recorded from DMS MSNs in mCherry- and hM4Di-injected 5xFAD mice. **C**, eEPSC amplitudes in DMS MSNs were smaller in hM4Di-injected 5xFAD mice than in mCherry-injected 5xFAD mice. Two-way RM ANOVA, *F*_(1,15)_ = 4.435, *p* = 0.052; Sidak’s multiple comparisons post-hoc analysis, \**p* < 0.05 versus hM4Di at the same stimulating intensities. n = 10 neurons from 4 mice (mCh); 7 neurons from 3 mice (hM4Di). **D**, PPR in DMS MSNs was higher in hM4Di-than in mCherry-injected 5xFAD mice. Two-way RM ANOVA, *F*_(1,17)_ = 5.213, *p* = 0.036. ^#^*p* < 0.05. n = 11 neurons from 4 mice (mCh); 8 neurons from 3 mice (hM4Di). **E**, Representative image showing red ACh sensor (rACh1.7) expression in striatal slices from 5xFAD mice. **F**, Sample traces of evoked striatal ACh release recorded from rACh1.7-expressing slices. **G and H**, The half-width (G), but not the amplitude (H), of evoked ACh release was greater in hM4Di-injected mice than in mCherry-injected 5xFAD controls. unpaired t-test, *t*_13_ = -2.448, *p* < 0.05 for half-width; Fig. 7D, *t*_13_ = 0.00147, *p* > 0.05 for amplitude. \**p* < 0.05. n = 8 slices from 4 animals (G and H, mCh), 7 slices from 4 animals (G and H, hM4Di). **I**, Sample traces of spontaneous striatal ACh release. **J-L**, The frequency (J), but not the amplitude (K) or half-width (L), of spontaneous ACh release was significantly higher in hM4Di-injected 5xFAD mice compared to mCherry-injected 5xFAD mice. unpaired t-test, (J), *t*_16_ = 2.377, *p* = 0.0303 for frequency; (K) *t*_16_ = 0.3087, *p* > 0.05 for amplitude; (L) *U* = 29, *p* > 0.05 for half-width. \**p* < 0.05. n = 9 slices from 4 animals (mCh and hM4Di).

**Supplementary Figure 10.**
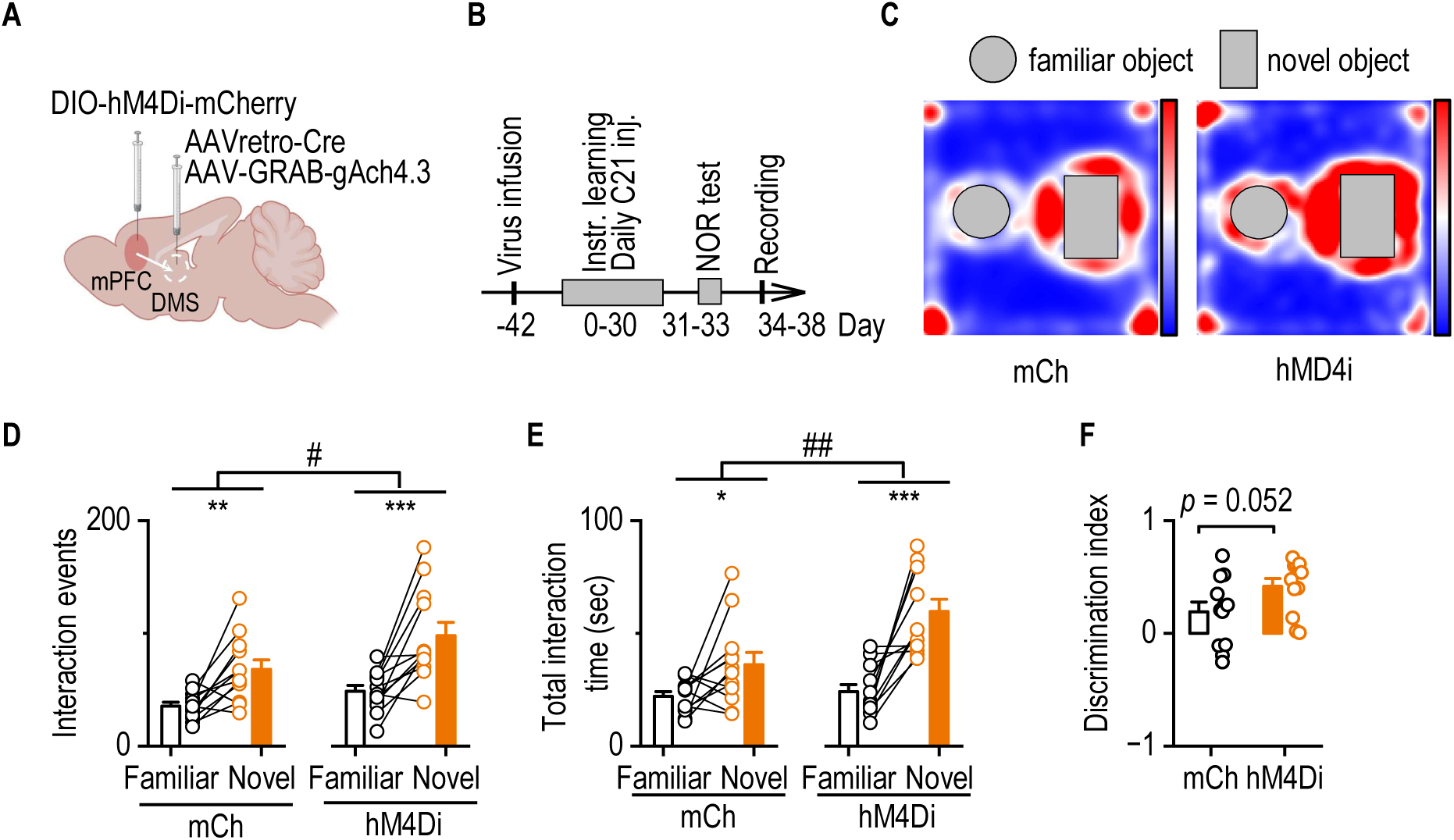
Sustained inhibition of mPFC to DMS circuit improved cognitive function in 5xFAD mice. **A,** Schematic illustrating viral infusion strategy in 5xFAD mice. AAV-retro-Cre and AAV-GRAB-ACh4m were infused into the DMS, and AAV-DIO-hM4Di-mCherry or AAV-DIO-mCherry into the mPFC. **B,** Experimental timeline. The novel object recognition (NOR) test was performed after one month of sustained chemogenetic inhibition. **C**, Heatmaps showing time spent interacting with novel and familiar objects during the 5-minute testing session. **D**, Both hM4Di- and mCherry-injected 5xFAD mice showed more interaction events with the novel object than the familiar object. However, hM4Di-injected mice exhibited significantly more novel object interactions than mCherry controls. Two-way RM ANOVA, *F*_(1,22)_ = 6.246, *p* = 0.020, ^#^*p* < 0.05; Sidak’s multiple comparisons post-hoc analysis, \**p* < 0.05. n = 12 mice (mCh and hM4Di). **E**, Both groups spent more total time interacting with the novel object than the familiar object, with hM4Di-injected mice exhibiting significantly longer interaction times overall. Two-way RM ANOVA, *F*_(1,22)_ = 10.753, *p* = 0.003, ^##^*p* < 0.01; Sidak’s multiple comparisons post-hoc analysis, \**p* < 0.05. n = 12 mice (mCh and hM4Di). **F**, hM4Di-injected 5xFAD mice showed a higher discrimination index than mCherry-injected 5xFAD mice. unpaired t-test, *t*_22_ = -2.055, *p* = 0.0519. n = 12 mice (mCh and hM4Di).

**Supplementary Figure 11.**
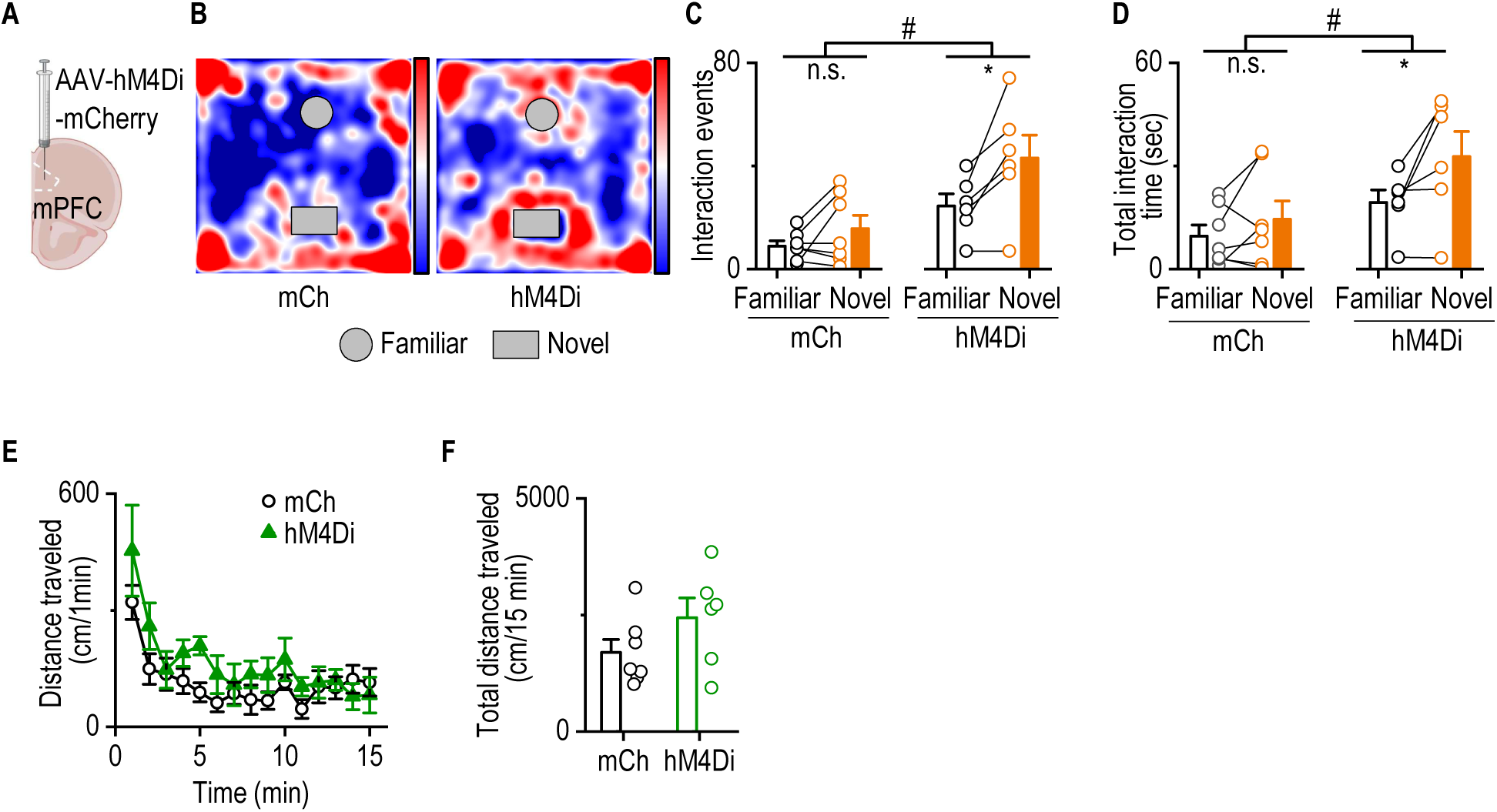
Sustained inhibition of mPFC neurons improves cognitive function in 5xFAD mice without impairing locomotor activity. **A**, Schematic showing AAV-hM4Di-mCherry or AAV-mCherry infusion in mPFC. **B**, Heatmaps of animals that interacted with novel and familiar objects. **C**, hM4Di-injected 5xFAD mice exhibited more interaction events with the novel object than controls in a 5-min session. Two-way RM ANOVA, *F*_(1,11)_ = 9.460, *p* = 0.011, ^#^*p* < 0.05; Sidak’s multiple comparisons post-hoc analysis, \**p* < 0.05. n = 7 mice (mCh); 6 mice (hM4Di). **D**, hM4Di-injected 5xFAD mice exhibited longer total interaction time with the novel object than controls in a 5-min session. Two-way RM ANOVA, *F*_(1,11)_ = 4.906, *p* = 0.049, ^#^*p* < 0.05; Sidak’s multiple comparisons post-hoc analysis, \**p* < 0.05. n = 7 mice (mCh); 6 mice (hM4Di). **E, F,** hM4Di-injected and mCherry-injected 5xFAD mice exhibited similar spontaneous locomotor activity as measured by time-course (E, Two-way RM ANOVA, *F*_(1,11)_ = 2.267, *p* > 0.05) and total distance traveled (F, unpaired t-test, *t*_11_ = -1.506, *p* > 0.05). n = 7 (E and F, mCh), 6 (E and F, hM4Di).

